# A Selective Multivalent NET-Associated Chromatin Neutralizer Resolves Infection-Associated Inflammation in Severe Sepsis

**DOI:** 10.64898/2026.06.29.735358

**Authors:** Chuanxu Cheng, Quanxin Ning, Juan Du, Jianati Dawulieti, Chenyang Guo, Madi Sun, Kunbao Zhang, Haijun Li, Qunjie Bi, Jian Li, Zhen Wu, Hanyao Huang, Zhi-Liang Ji, Jin-Zhi Du, Chao Yang, Dan Shao, Kam W. Leong

## Abstract

Targeting the overwhelming inflammation driven by neutrophil extracellular traps (NETs) during infection provides an opportunity to manage severe sepsis. This potential needs to be realized by exploring selective NET-neutralization materials, which remains a challenge. Herein, we report a multivalent macromolecular strategy that targets NET-associated DNA-histone chromatin complexes while preserving antibacterial activity of aminoglycoside. We identify 8-arm PEG-conjugated netilmicin (8-arm Netil) as a lead NETs-neutralizer from a library of multivalent aminoglycoside-displayed materials. When compared with 2- and 4-arm counterparts, 8-arm Netil exhibits potent antibacterial activity and high-affinity binding to DNA-histone chromatin complexes through stable multivalent noncovalent interactions, thereby suppressing NET-induced TLR4/TLR9 activation and macrophage inflammatory responses. In severe septic mice, intravenously administered 8-arm Netil preferentially accumulates in inflamed tissues, leading to improved survival protection, owing to the reduction of bacterial dissemination, NET accumulation, systemic cytokine production, and multiple-organ injury. These findings establish NET-associated DNA-histone chromatin complexes as actionable extracellular targets and demonstrate multivalent chromatin targeting as a rational material design strategy for selective NET neutralization and inflammation control in severe sepsis.

## 1. Introduction

Sepsis is a life-threatening organ dysfunction caused by a dysregulated host response to infection and remains a major global health challenge despite advances in antimicrobial therapy, source control, and intensive care management^[1–3]^. Although microbial infection initiates sepsis, disease progression is not determined solely by pathogen burden^[4–5]^. During severe sepsis, bacterial products, host-derived damage signals, and activated immune cells reinforce each other to drive systemic cytokine release, vascular dysfunction, immune imbalance, and multiple-organ injury^[6–8]^. Antibiotics are indispensable for infection control^[1]^, yet bacterial killing alone does not eliminate the inflammatory molecular debris generated during infection, tissue injury, and antimicrobial-mediated bacterial disruption^[9–10]^. Conversely, broad anti-inflammatory intervention may suppress harmful inflammation but can compromise host defense^[6–7, 11]^. Therefore, an effective therapeutic strategy for severe sepsis should coordinate pathogen control with selective removal of inflammatory signals that sustain immune dysregulation.

Neutrophil extracellular traps (NETs) are a critical interface between antimicrobial defense and inflammatory injury in sepsis^[12–14]^. NETs are extracellular chromatin networks composed of DNA-histone scaffolds decorated with neutrophil-derived enzymes and antimicrobial proteins^[12–13]^. Although NETs can entrap and restrict microorganisms, excessive NET formation or impaired NET clearance converts this protective response into a persistent inflammatory stimulus^[12–16]^. In particular, the NET-associated DNA-histone chromatin complex serves as both a structural scaffold and a biochemical source of inflammatory activation^[17–18]^. NET-associated DNA, histones, and chromatin-associated proteins can engage innate immune pathways, including Toll-like receptor 4 (TLR4)- and Toll-like receptor 9 (TLR9)-mediated NF-κB signaling, thereby amplifying cytokine production, tissue damage, and organ dysfunction^[10, 18–21]^. Thus, the DNA-histone chromatin complex within NETs is not merely a marker of neutrophil activation, but a druggable extracellular target linking infection, NET accumulation, and non-resolving inflammation.

Current NET-targeting strategies remain insufficient for severe sepsis. DNase-mediated degradation can digest extracellular DNA, but it does not fully neutralize histones, chromatin-associated proteins, or other NET-derived inflammatory components^[22, 23]^. DNA fragmentation may also leave residual bioactive complexes, whereas direct inhibition of NET formation may impair neutrophil-mediated antimicrobial defense during active infection^[12, 20, 24]^. These limitations suggest that selectively binding and neutralizing intact NET-associated DNA-histone chromatin complexes may be more effective than simply degrading NET DNA or broadly suppressing neutrophil function. In contrast to cationic nucleic-acid scavenging nanomaterials that bind cell-free DNA non-selectively, the strategy pursued here is designed to recognize the assembled DNA-histone chromatin complex rather than free DNA alone, which we hypothesized would improve selectivity toward NET-associated inflammatory scaffolds. The extent of this selectivity is quantified in **Section 2.3**. Ideally, such a therapeutic system should reduce pre-existing extracellular chromatin burden, suppress NET-induced inflammatory signaling, and retain antibacterial activity to limit bacteria-driven NET formation. Multivalent biomaterials offer a rational design principle to meet these requirements^[25–28]^. Biological recognition events involving chromatin, nucleic acids, and protein complexes are often spatially organized and avidity-driven^[27, 29–30]^. We therefore hypothesized that presenting multiple chromatin-interacting ligands on a macromolecular scaffold could convert weak monovalent interactions into high-avidity binding toward NET-associated DNA-histone chromatin complexes^[25–28, 31]^. Aminoglycosides were selected as functional ligands because they possess intrinsic antibacterial activity and contain cationic, hydrogen-bonding-rich structures capable of interacting with nucleic acid-protein assemblies^[32–35]^. Conjugating aminoglycosides onto multi-arm polyethylene glycol (PEG) could therefore generate a dual-function polymer antibiotic: the aminoglycoside motifs would preserve antibacterial activity, whereas the multivalent PEG architecture would enable selective extracellular chromatin binding and NET scavenging. By increasing the number of aminoglycoside ligands displayed on the PEG, this strategy was designed to strengthen multivalent chromatin binding while retaining antibacterial activity, thereby integrating infection control with NET-associated chromatin neutralization and inflammatory resolution.

Here, we report a multivalent macromolecular aminoglycoside platform for severe sepsis therapy and identify 8-arm PEG-conjugated netilmicin (8-arm Netil) as a lead NET-associated chromatin-neutralizing material. By screening free aminoglycosides and their 2-, 4-, and 8-arm PEG conjugates, we found that 8-arm Netil combined strong DNA-histone chromatin complex binding, NETs scavenging, antibacterial activity, biocompatibility, and in vivo anti-sepsis efficacy. Mechanistic studies revealed that 8-arm Netil forms stable multivalent noncovalent interactions with DNA-histone chromatin complexes, thereby reducing NETs-induced TLR4/TLR9 activation and macrophage inflammatory responses. In *Escherichia coli*-induced bacterial peritonitis and cecal ligation and puncture (CLP)-induced severe sepsis models, intravenously administered 8-arm Netil reduced bacterial dissemination, NETs accumulation, systemic cytokine production, and multiple-organ injury, leading to markedly improved survival. Flow cytometry, transcriptomic analysis, and tissue immunostaining further showed that 8-arm Netil reshaped immune-cell composition and suppressed inflammatory signaling in blood, peritoneal fluid, and intestinal tissues. Together with favorable pharmacokinetic behavior, inflamed-tissue accumulation, hemocompatibility, and systemic biosafety, these findings identify NET-associated DNA-histone chromatin complexes as actionable extracellular targets and demonstrate that multivalent ligand display can convert a conventional antibiotic into a selective NET-neutralizing material, integrating infection control with inflammation resolution in severe sepsis.

## 2. Results and Discussion

### 2.1. Construction and characterization of multivalent PEGylated aminoglycosides

To develop a multivalent platform for chromatin-targeting aminoglycosides (AG), we synthesized a series of PEGylated aminoglycoside conjugates in which the number of displayed aminoglycoside ligands was controlled using 2-, 4-, and 8-arm PEG. The synthetic strategy involved two sequential steps: terminal carboxylation of 2-, 4-, or 8-arm PEG-OH through reaction with suberic anhydride, followed by amide coupling between PEG-COOH and aminoglycoside ligands (**Figure 1a, b**). This modular route enabled the generation of aminoglycoside conjugates with different ligand densities while preserving the same PEG-based macromolecular framework. Eleven aminoglycosides, including netilmicin (Netil), amikacin (Ami), ribostamycin (Rib), sisomicin (Sis), paromomycin (Par), streptomycin (Str), kanamycin A (Kan), tobramycin (Tob), gentamicin (Gen), apramycin (Apr), and neomycin (Neo) were selected as ligand candidates (**Figure 1b**).

The successful synthesis of 8-arm PEG-conjugated netilmicin (8-arm Netil) was then verified as a representative lead construct. FT-IR spectroscopy confirmed the formation of amide bonds between 8-arm PEG-COOH and netilmicin, as evidenced by the disappearance of the carboxylic acid C=O stretching band at around 1733 cm^-1^ and the emergence of amide-associated absorption bands after conjugation (**Figure 1d**). ^1^H NMR spectroscopy further supported successful netilmicin grafting onto the 8-arm PEG. The final 8-arm Netil conjugate displayed characteristic proton signals from the PEG-suberate linker and netilmicin, and the integration ratio between the assigned linker protons and netilmicin protons was consistent with near-complete substitution of the eight terminal carboxyl groups by netilmicin (**Figure 1c**). GPC analysis further corroborated this controlled conjugation, showing the expected molecular-weight increase and a relatively narrow molecular-weight distribution compared with 8-arm PEG-OH (**Figure 1e**). These combined structural and molecular-weight analyses demonstrate that 8-arm Netil was prepared as a well-defined multivalent aminoglycoside conjugate with controlled ligand valency.

**Figure 1.**
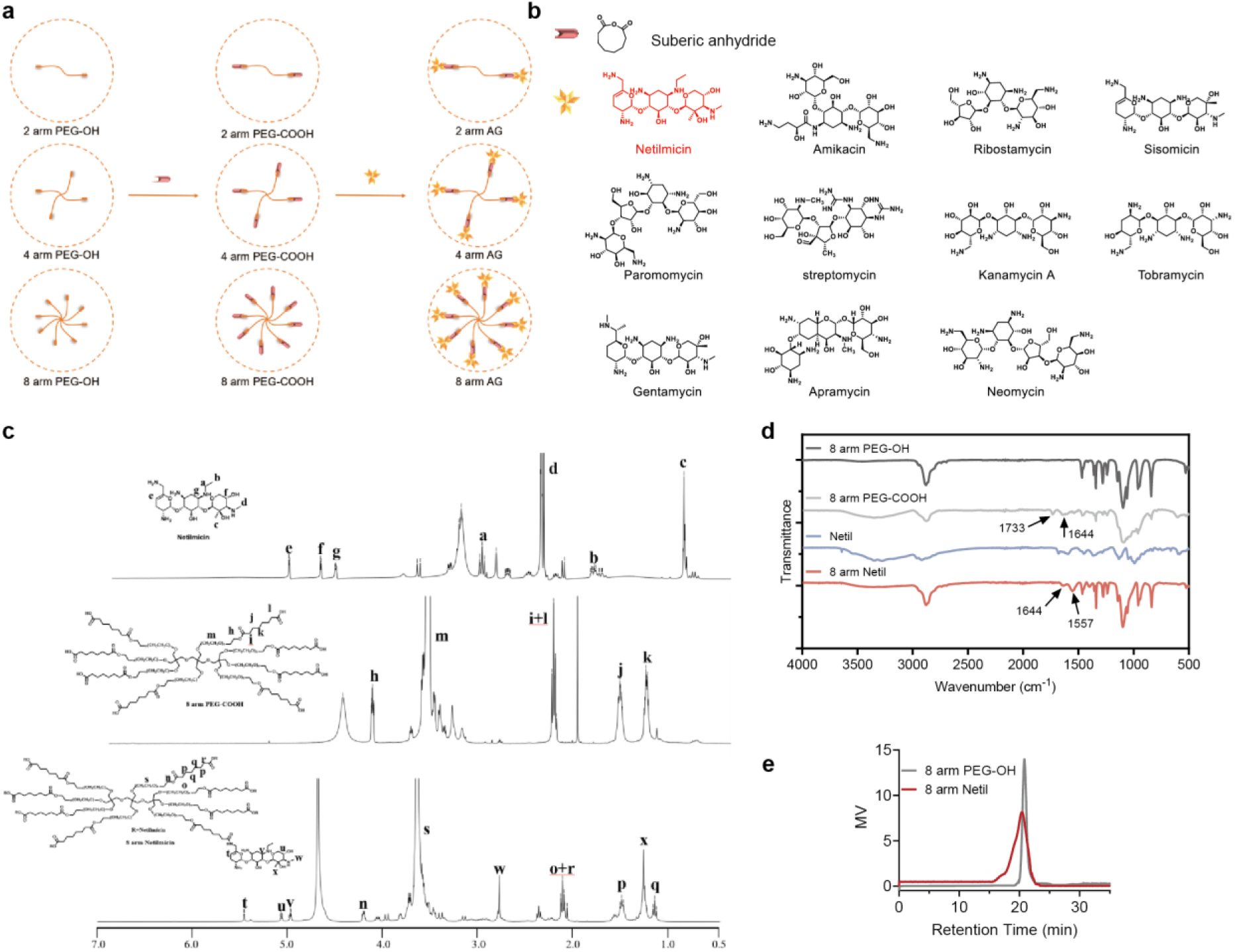
Synthesis and characterization of multivalent 8-arm aminoglycoside conjugates. **(a**) Schematic illustration of the synthetic route for 2-, 4-, and 8-arm aminoglycoside (AG) conjugates. Multi-arm PEG-OH was first converted to PEG-COOH using suberic anhydride and subsequently conjugated with aminoglycosides to generate multivalent PEG-AG. (**b**) Chemical structures of suberic anhydride and the aminoglycosides used in this study, including netilmicin (Netil), amikacin (Ami), ribostamycin (Rib), sisomicin (Sis), paromomycin (Par), streptomycin (Str), kanamycin A (Kan), tobramycin (Tob), gentamicin (Gen), apramycin (Apr), and neomycin (Neo). (**c**) ^1^H NMR spectra of Netil, 8-arm PEG-COOH, and 8-arm Netil in DMSO-d6. (**d**) FT-IR spectra of 8-arm PEG-OH, 8-arm PEG-COOH, Netil, and 8-arm Netil. **(e**) Gel permeation chromatography (GPC) traces of 8-arm PEG-OH and 8-arm Netil.

### 2.2 Screening identifies 8-arm Netil as a lead NET-associated chromatin neutralizer

After establishing the multivalent PEGylated aminoglycoside library, we next screened these conjugates to identify candidates capable of coordinating antibacterial activity with extracellular chromatin scavenging. Because NETs are composed primarily of DNA-histone chromatin complexes, we first evaluated the capacity of free aminoglycosides and their 2-, 4-, and 8-arm PEG conjugates to bind extracellular chromatin. 8-arm aminoglycoside conjugates showed markedly higher binding than their free, 2-arm, or 4-arm counterparts (**Figure 2a** and **Figure S1**), showing a strong valency-dependent enhancement in chromatin-binding efficiency. Among them, 8-arm Netil displayed one of the strongest extracellular chromatin-binding capacities. Importantly, this enhanced chromatin interaction was not accompanied by obvious cytotoxicity or hemocompatibility concerns, as most 8-arm aminoglycoside conjugates showed minimal hemolysis and maintained high cell viability even at elevated concentrations (640 μg/mL) **(Figure S2)**.

We then asked whether the chromatin-binding capacity of these macromolecular aminoglycosides translated into therapeutic efficacy in vivo. Using a severe cecal ligation and puncture (CLP)-induced sepsis model, one of the gold standards in studying sepsis^[36]^, mice were intravenously treated with different 8-arm aminoglycoside conjugates at 1, 12, and 24 h after CLP (**Figure 2b**). Unfortunately, 90% of CLP-induced animals died within 72 h without any treatment. In contrast, 6/11 macromolecular AGs treatments resulted in a more than 60% survival rate, with increased body weight (**Figure 2c** and **Figure S3**). Notably, 8-arm Netil provided the strongest protection, achieving 100% survival rate under this treatment setting; the untreated control reached approximately 90% mortality within 72 h, confirming a lethal challenge under the puncture protocol used. Body-weight monitoring further supported the superior therapeutic benefit of selected 8-arm conjugates compared with untreated CLP mice or free aminoglycoside controls **(Figure S5**).

Because aminoglycosides are antibacterial agents, we next examined whether antibacterial potency alone explained the anti-sepsis efficacy of the polymer conjugates. The minimum inhibitory concentration (MIC) and minimal bactericidal concentration (MBC) assays against *Escherichia coli* showed that PEGylated aminoglycosides generally retained antibacterial activity, and in several cases showed improved antibacterial performance compared with free aminoglycosides (**Figure 2d, e**). However, survival after 8-arm AG treatment showed little association with antibacterial potency, as indicated by weak correlations with MIC and MBC values against *E. coli* (MIC: Pearson r = 0.079, n = 11, P = 0.8166; MBC: Pearson r = 0.1168, n = 11, P = 0.732; **Figure 2g, h**), with the corresponding 95% confidence intervals spanning zero. In contrast, survival after free AG treatment showed a stronger inverse association with MIC and MBC values, indicating that lower MIC/MBC values were associated with higher survival (MIC: Pearson r = −0.813, n = 11, P = 0.0023; MBC: Pearson r = −0.787, n = 11, P = 0.0040; **Figure 2i** and **Figure S4**). Because these free-AG analyses were based on a limited compound set and included overlapping antibacterial values, they are reported with exact P values and 95% confidence intervals in the figure legend and are interpreted as exploratory. Notably, extracellular chromatin neutralization by 8-arm AGs showed a stronger positive correlation with survival (Pearson r = 0.7568, n = 11, P = 0.0070; **Figure 2f**). A second, complementary readout based on NET/chromatin signal neutralization in septic serum showed a consistent positive association with survival (**Figure 2o**, Pearson r = 0.8802). These two assays measure distinct chromatin-associated activities—neutralization of reconstituted DNA**-**histone chromatin complexes in vitro and suppression of NET/chromatin signals in septic serum—but their concordant direction supports a link between chromatin neutralization and therapeutic benefit. Together, these results suggest that the anti-sepsis activity of macromolecular aminoglycosides is not governed by antibacterial potency alone, but is closely associated with their ability to neutralize NET-associated chromatin.

In addition to infection, severe sepsis is driven by systemic inflammatory response syndrome (SIRS), which involves the dysregulation of multiple danger signals^[8]^. We therefore compared the direct neutralization of representative sepsis-associated inflammatory mediators and danger signals, including LPS, HMGB-1, TNF-α, and extracellular chromatin. *In vitro* screening showed that 8-arm AG preferentially neutralized extracellular chromatin rather than soluble inflammatory mediators (**Figure 2j** and **Figure S8**). This selectivity was further confirmed using serum from CLP-induced septic mice and patients with sepsis, where the most pronounced reduction was consistently observed for chromatin/NET-associated signals, rather than LPS, HMGB-1, or TNF-α (**Figure 2k, Figures S6** and **S7**). Correlation analysis further supported this conclusion: survival showed weak associations with LPS, TNF-α, and HMGB-1 binding efficiencies, with correlation coefficients of 0.027, 0.009, and 0.095, respectively (**Figure 2l-n**), but strongly correlated with NET/chromatin binding efficiency (r = 0.8802; **Figure 2o**). Natural NET-binding assays using neutrophil-derived NETs provided additional evidence that 8-arm AG, particularly 8-arm Netil, directly interact with NET-associated chromatin **(Figure S8e, f**).

Among the screened candidates, 8-arm Netil showed the most favorable combination of extracellular chromatin binding, NET neutralization, antibacterial activity, biocompatibility, and in vivo anti-sepsis efficacy. Therefore, we identified 8 arm Netil as our lead macromolecular AG candidate from the library and screening tests for further studies.

**Figure 2.**
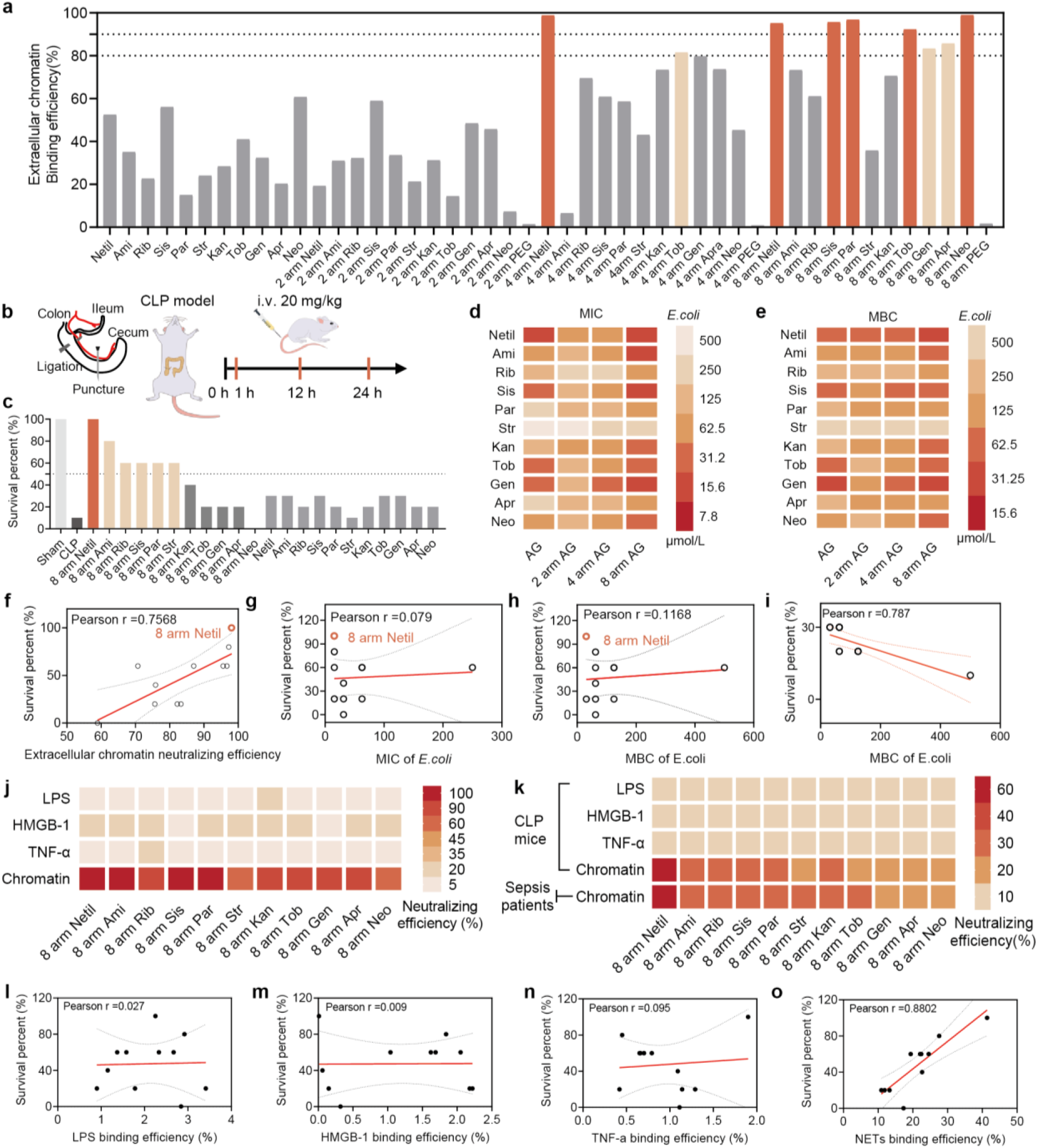
*In vitro* and *in vivo* screening of a macromolecular aminoglycoside library identifies 8-arm Netil as a lead candidate for sepsis therapy. (a) Extracellular chromatin-binding efficiencies of free aminoglycosides and their corresponding 2-, 4-, and 8-arm PEG conjugates at a material-to-chromatin mass ratio of 1:1. **(b**) Schematic illustration of the severe cecal ligation and puncture (CLP) model and treatment schedule. BALB/c mice received intravenous injections of aminoglycoside conjugates or free aminoglycosides at 1, 12, and 24 h after CLP. **(c**) Survival percentages of CLP-challenged mice treated with 8-arm aminoglycoside conjugates or corresponding free aminoglycosides. **(d, e**) Heatmaps showing the minimum inhibitory concentration (MIC; **d**) and minimum bactericidal concentration (MBC; **e**) of free aminoglycosides and their 2-, 4-, and 8-arm conjugates against *E. coli* (ATCC 25922). **(f-i**) Correlation analyses between survival percentage and extracellular chromatin-binding efficiency **(f**), MIC of *E. coli* for 8-arm aminoglycoside conjugates **(g**), MBC of *E. coli* for 8-arm aminoglycoside conjugates **(h**), and MBC of *E. coli* for free aminoglycosides **(i**). For **f,** Pearson r = 0.7568, n = 11, P = 0.0070, and the 95% confidence interval (CI) for the regression slope was 0.6223 to 2.945. For **g,** Pearson r = 0.079, n = 11, P = 0.8166, and the 95% CI for the regression slope was −0.2974 to 0.3676. For **h,** Pearson r = 0.1168, n = 11, P = 0.7323, and the 95% CI for the regression slope was −0.1429 to 0.1958. For **i,** Pearson r = −0.787, n = 11, P = 0.0040, and the 95% CI for the regression slope was −0.06358 to −0.01634. **(j**) Scavenging efficiencies of 8-arm aminoglycoside conjugates toward inflammatory danger signals, including LPS, HMGB-1, TNF-α, and chromatin *in vitro*. **(k**) Scavenging efficiencies of 8-arm aminoglycoside conjugates toward inflammatory danger signals in serum from CLP-induced septic mice and patients with sepsis. **(l-o**) Correlation analyses between survival percentage and binding efficiencies toward LPS **(l),** HMGB-1 **(m),** TNF-α **(n),** and NETs **(o**) in serum from CLP-induced septic mice. For **l,** Pearson r = 0.027, n = 11, P = 0.9363, and the 95% CI for the regression slope was −26.88 to 28.91. For **m,** Pearson r = 0.009, n = 11, P = 0.9782, and the 95% CI for the regression slope was −24.74 to 25.37. For **n,** Pearson r = 0.095, n = 11, P = 0.7807, and the 95% CI for the regression slope was −45.31 to 58.48. For **o,** Pearson r = 0.8802, n = 11, P = 0.0005, and the 95% CI for the regression slope was 1.743 to 4.345. Data are presented as mean values. n = 3 independent experiments for in vitro assays, n = 10 mice per group for survival studies, and n = 11 compounds or conjugates for correlation analyses unless otherwise indicated.

### 2.3 8-arm Netil selectively recognizes DNA-histone chromatin complexes through stable multivalent noncovalent interactions

Having identified 8-arm Netil as the lead candidate, we next investigated the molecular basis underlying its selective recognition of NET-associated DNA-histone chromatin complexes. Molecular docking suggested that both free Netil and 8-arm Netil could engage the DNA-histone chromatin complex at phosphate-rich and protein-associated regions, but the multivalent architecture of 8-arm Netil enabled simultaneous interactions across a broader chromatin interface (**Figure 3a, b**). To further assess binding stability, molecular dynamics simulations were performed using free Netil and 2-, 4-, and 8-arm Netil. During the 50 ns simulation, free Netil showed limited and unstable association with the chromatin complex, whereas chromatin retention progressively increased with the number of PEG arms (**Figure 3c** and **Figure S9**). Binding interaction-energy analysis further supported this valency-dependent enhancement, with calculated interaction energies (estimated by the molecular mechanics/Poisson–Boltzmann surface area approach, which neglects configurational entropy and therefore provides relative free energies) decreasing from −17.753 kJ/mol for free Netil to −80.709, −92.341, and −203.409 kJ/mol for 2-, 4-, and 8-arm Netil, respectively (**Figure 3c**). These results demonstrate that increasing the number of PEG arms enhances Netil-chromatin binding stability, with 8-arm Netil forming the most stable multivalent complex with the DNA-histone chromatin structure.

**Figure 3.**
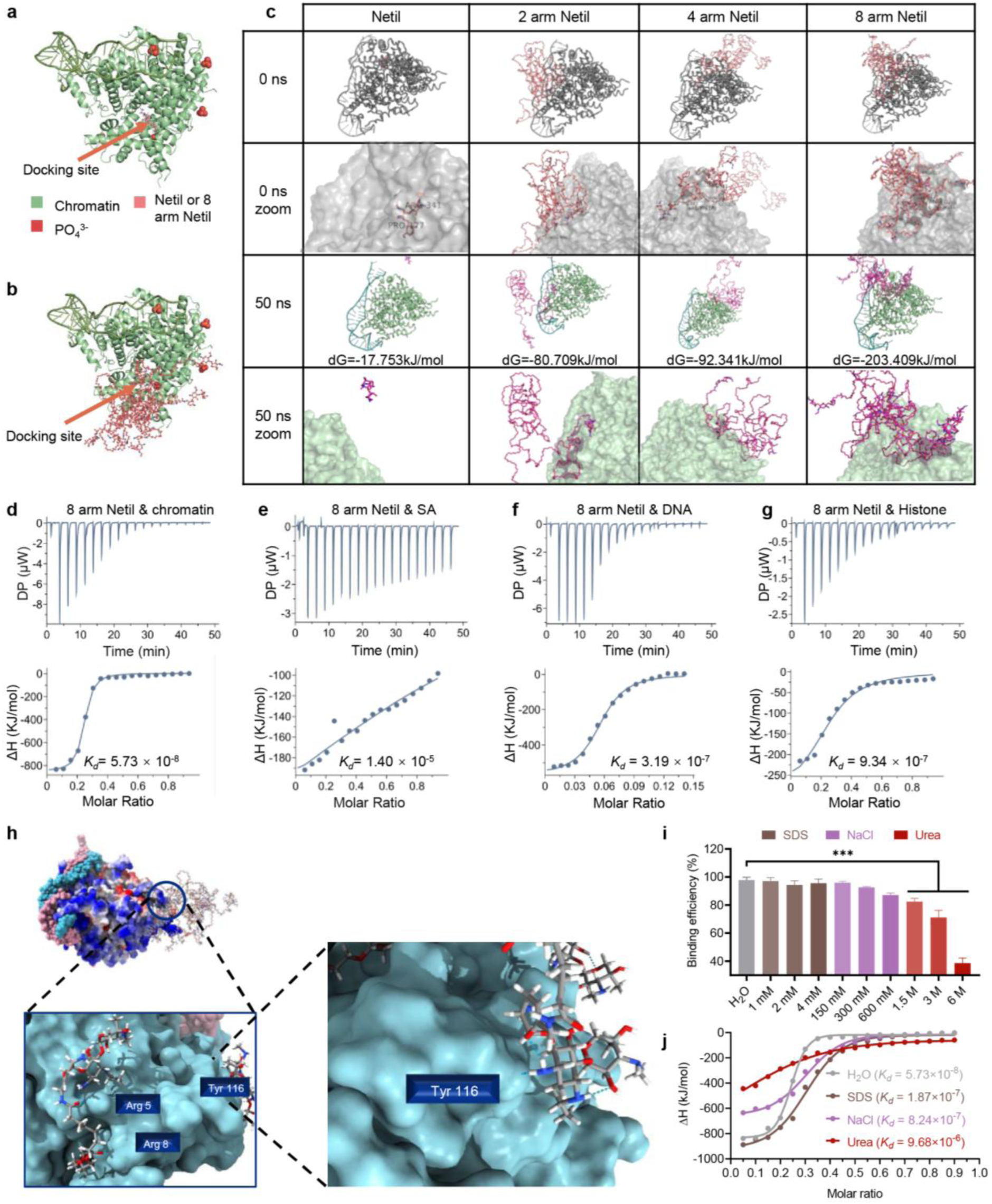
Molecular and thermodynamic basis of selective NET binding by 8-arm Netil. **(a, b)** Molecular docking models showing the interactions of free Netil **(a)** and 8-arm Netil **(b)** with a DNA/histone chromatin complex. Chromatin is shown in green, phosphate groups are highlighted in red, and Netil or 8-arm Netil is shown in pink. **(c)** Molecular dynamics simulations comparing the binding stability of free Netil and 2-, 4-, and 8-arm Netil with chromatin over 50 ns. Representative snapshots at 0 and 50 ns, together with magnified views of the binding interfaces, are shown, with calculated interaction energies indicated. **(d-g)** Isothermal titration calorimetry (ITC) analysis of 8-arm Netil binding to the DNA/histone chromatin complex **(d)**, serum albumin (SA; **e)**, DNA **(f)**, and histone **(g)**. The dissociation constant (Kd) was obtained by fitting the integrated heat release as a function of molar ratio. **(h)** Molecular docking analysis showing the local interaction interface between 8-arm Netil and the DNA/histone chromatin complex, highlighting representative binding residues and contact sites. **(i)** Binding efficiencies of 8-arm Netil toward the DNA/histone chromatin complex in the presence of sodium dodecyl sulfate (SDS), sodium chloride (NaCl), or urea. SDS was used to perturb hydrophobic interactions, NaCl to screen electrostatic interactions, and urea to disrupt hydrogen-bonding interactions. **(j)** ITC analysis of 8-arm Netil binding to the DNA/histone chromatin complex in H₂O or in the presence of SDS, NaCl, or urea, revealing the contribution of noncovalent interactions to chromatin/NET recognition. Data are presented as mean ± s.e.m.; n = 3 independent experiments. ITC measurements and molecular dynamics simulations were independently repeated three times. ΔH, enthalpy change. Statistical significance was calculated by one-way ANOVA with Tukey’s multiple-comparison test; ***P < 0.001.

We next quantified the binding affinity and selectivity of 8-arm Netil using isothermal titration calorimetry (ITC). 8-arm Netil bound the DNA-histone chromatin complex with high affinity, with a dissociation constant (*Kd*) of 5.73 × 10^-8^ M (**Figure 3d)**. By contrast, its affinity for serum albumin was much weaker, with a *Kd* of 1.40 × 10^-5^ M, supporting preferential chromatin recognition over abundant serum proteins during systemic administration (**Figure 3e)**. This corresponds to an approximately 240-fold preference for the chromatin complex over serum albumin, an abundant and clinically relevant off-target protein during intravenous administration. Among the tested off-target substrates, serum albumin showed the largest selectivity margin relative to the DNA–histone chromatin complex, supporting preferential chromatin recognition by 8-arm Netil under systemic delivery conditions. 8-arm Netil also showed lower affinity for isolated DNA and histone than for the intact DNA-histone chromatin complex, with *Kd* values of 3.19 × 10^-7^ M and 9.34 × 10^-7^ M, respectively (**Figure 3f, g**). These results indicate that 8-arm Netil binds with modestly higher affinity to the assembled DNA-histone chromatin complex (approximately 5.6-fold and 16-fold tighter than to isolated DNA or histone, respectively). Consistently, ITC analysis of additional Netil conjugates and control cationic polymers showed that efficient and selective chromatin recognition requires both the chromatin-interacting Netil ligand and its multivalent display on the PEG **(Figure S10**). Although fourth-generation poly(amidoamine) dendrimer (PAMAM G4), a classical nucleic acid-binding cationic polymer^[37]^, showed measurable chromatin-binding affinity, its lower selectivity highlights the importance of the 8-arm Netil architecture for preferential NET-associated chromatin scavenging **(Figure S10**). Together, these ITC results demonstrate that 8-arm Netil achieves high-affinity and selective recognition of intact DNA-histone chromatin complexes through multivalent ligand presentation.

To clarify the interaction mode, we analyzed the local binding interface between 8-arm Netil and the DNA-histone chromatin complex. Docking visualization showed that 8-arm Netil established multiple contact sites around chromatin-associated residues, including Arg5, Arg8, and Tyr116, consistent with a multivalent binding mode involving electrostatic, hydrogen-bonding, and protein-associated interactions (**Figure 3h**). Competitive binding assays further confirmed the contribution of these noncovalent forces. Chromatin-binding efficiency remained relatively stable under SDS treatment and moderate ionic strength but was progressively reduced by increasing urea concentrations, indicating a substantial contribution of hydrogen-bonding interactions to stable complex formation (**Figure 3i** and **Figure S11**). Consistently, ITC measurements showed that SDS, NaCl, and urea weakened the interaction between 8-arm Netil and the DNA-histone chromatin complex, increasing the *Kd* from 5.73 × 10^-8^ M in water to 1.87 × 10^-7^ M, 8.24 × 10^-7^ M, and 9.68 × 10^-6^ M, respectively (**Figure 3j**). Together, these results demonstrate that 8-arm Netil recognizes NET-associated DNA-histone chromatin complexes through stable, high-avidity, multivalent noncovalent interactions, with hydrogen bonding making a major contribution to complex stabilization.

### 2.4 8-arm Netil binds NET-associated chromatin and neutralizes NET-driven inflammatory activation

After establishing the high-affinity interaction between 8-arm Netil and DNA-histone chromatin complexes, we next examined whether this binding translated into direct NET recognition and inflammatory neutralization. To model the major chromatin scaffold of NETs, DNA and histones were assembled into reconstituted DNA-histone chromatin complexes (**Figure 4a**). In parallel, native NETs (n-NETs) were collected from PMA-stimulated neutrophils^[38]^ (**Figure 4b**). Scanning electron microscopy confirmed that both reconstituted chromatin complexes and n-NETs formed web-like extracellular networks with nanoscale-to-microscale architectures resembling NET-associated chromatin structures (**Figure 4c, d**). Fluorescence imaging showed that PEGylation with increasing arm numbers improved the binding of Netil to chromatin/NET structures. Among all Netil formulations, 8-arm Netil showed the strongest colocalization with DNA-histone chromatin complexes, with the material fluorescence mainly overlapping with DNA- and histone-positive chromatin regions (**Figure 4e**). A similar trend was observed for n-NETs, where 8-arm Netil displayed the highest colocalization with extracellular NET structures marked by DNA and citrullinated histone H3 (Cit H3) (**Figure 4f**). These results confirm that the intact DNA-histone chromatin complex is a major binding substrate within NETs and that the 8-arm architecture enables efficient recognition of native extracellular chromatin networks. In RAW264.7 macrophages, internalized n-NET-associated 8-arm Netil signals partially colocalized with lysosomal compartments, suggesting that 8-arm Netil-mediated NET complexation may facilitate intracellular processing of NET-associated chromatin after macrophage uptake (**Figure 4g**).

We then assessed whether 8-arm Netil-mediated NET binding and neutralizing could suppress inflammatory signaling. First, control reporter assays confirmed that free Netil and Netil-based PEG conjugates did not directly activate HEK-Blue TLR4 or TLR9 reporter cells, excluding intrinsic TLR agonism of the materials themselves **(Figure S12)**. In contrast, n-NETs strongly activated TLR4 and TLR9 reporter cells and induced TNF-α secretion from RAW264.7 macrophages (**Figure 4h-j**). DNase I efficiently reduced TLR9 activation and macrophage TNF-α production, but showed limited suppression of TLR4 activation, consistent with incomplete neutralization of non-DNA NET-associated inflammatory components. By comparison, 8-arm Netil markedly reduced n-NET-induced TLR4 activation, TLR9 activation, and macrophage inflammatory responses, outperforming free Netil and lower-valency Netil conjugates (**Figure 4h-j**). To test whether this anti-inflammatory effect extended to complex biological samples, we evaluated inflammatory activation induced by serum from CLP-induced septic mice and patients with sepsis. Both CLP mouse serum and sepsis patient serum activated TLR4/TLR9 signaling and stimulated macrophage TNF-α release, reflecting the presence of circulating inflammatory danger signals, including NET-associated chromatin. Treatment with 8-arm Netil substantially suppressed TLR4 and TLR9 activation and reduced TNF-α production in these serum-stimulation models, with stronger effects than free Netil or lower-valency conjugates (**Figure 4k-p**). Dose-response experiments further confirmed that 8-arm Netil inhibited n-NET-, CLP serum-, and patient serum-induced inflammatory activation in a concentration-dependent manner **(Figure S13**).

Finally, we asked whether NETs retained inflammatory activity after direct complexation with 8-arm Netil. Native NETs were preincubated with free Netil or Netil-based PEG conjugates, and the resulting NET/material complexes were then used to stimulate TLR reporter cells and macrophages. Compared with untreated NETs or complexes formed with lower-valency conjugates, NET/8-arm Netil complexes showed markedly reduced activation of TLR4 and TLR9 reporter cells and induced substantially lower TNF-α production in macrophages **(Figure S14)**. These results indicate that 8-arm Netil does not simply colocalize with NETs or mask fluorescence signals, but functionally neutralizes the inflammatory activity of NET-associated chromatin. Together, these findings demonstrate that 8-arm Netil selectively binds DNA-histone chromatin complexes within native NETs and suppresses NET-driven inflammatory activation through multivalent extracellular chromatin complexation.

**Figure 4.**
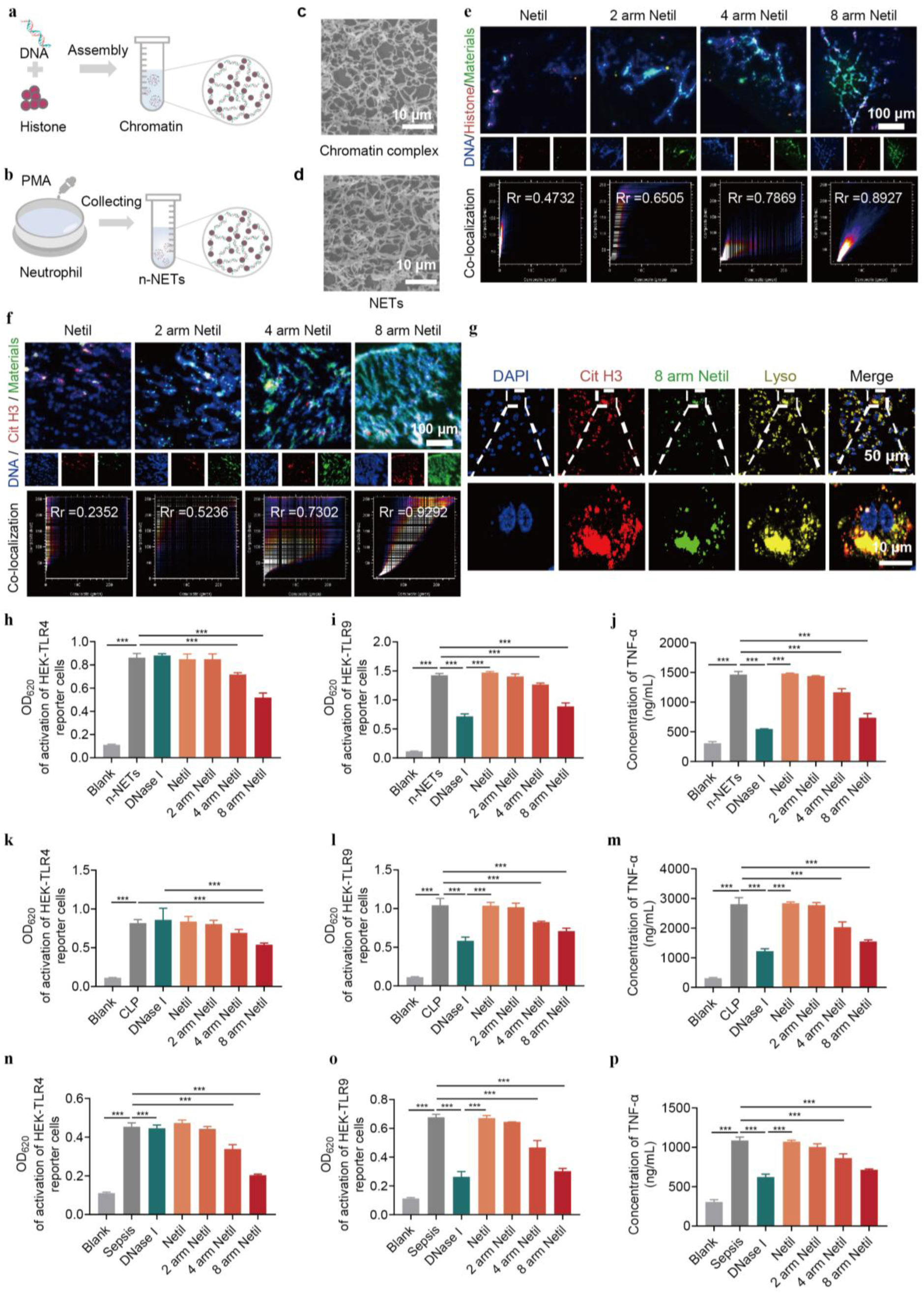
8-arm Netil selectively binds chromatin/NETs and suppresses NETs-associated inflammatory activation. **(a)** Schematic illustration of the preparation of a DNA**/**histone chromatin complex by self-assembly of DNA and histone. **(b)** Schematic illustration of the induction and collection of natural NETs (n-NETs) from PMA-stimulated neutrophils. **(c, d)** Scanning electron microscopy images of the DNA**/**histone chromatin complex **(c)** and n-NETs **(d)**. Scale bars, 10 μm. **(e, f)** Confocal fluorescence images and colocalization analysis showing the binding of free Netil and 2-, 4-, and 8-arm Netil to the DNA**/**histone chromatin complex **(e)** and n-NETs **(f)**. DNA was stained with DAPI, histone or citrullinated histone H3 (Cit H3) was labeled in red, and aminoglycoside materials were labeled in green. Scale bars, 100 μm. **(g)** Confocal fluorescence images showing the intracellular colocalization of n-NETs and 8-arm Netil with lysosomes in RAW264.7 macrophages after incubation. DNA, Cit H3, 8-arm Netil, and lysosomes are shown in blue, red, green, and yellow, respectively. Scale bars, 50 μm and 10 μm for magnified images. **(h-j)** Activation of HEK-TLR4 reporter cells **(h)**, HEK-TLR9 reporter cells **(i)**, and RAW264.7 macrophages **(j)** after stimulation with n-NETs in the presence of DNase I, free Netil, or 2-, 4-, and 8-arm Netil. **(k-m)** Activation of HEK-TLR4 reporter cells **(k)**, HEK-TLR9 reporter cells **(l)**, and RAW264.7 macrophages **(m)** after stimulation with serum from CLP-induced septic mice in the presence of DNase I, free Netil, or 2-, 4-, and 8-arm Netil. **(n-p)** Activation of HEK-TLR4 reporter cells **(n)**, HEK-TLR9 reporter cells **(o)**, and RAW264.7 macrophages **(p)** after stimulation with serum from septic patients in the presence of DNase I, free Netil, or 2-, 4-, and 8-arm Netil. Data are presented as mean ± s.e.m. n = 4 biologically independent samples in **h-p**. Statistical significance was calculated by one-way ANOVA with Tukey’s multiple-comparison test; ***P < 0.001.

### 2.5 8-arm Netil provides survival protection in bacterial peritonitis

To determine whether 8-arm Netil could coordinate antibacterial activity with NET-associated inflammatory control during active infection, we first evaluated its therapeutic efficacy in an *Escherichia coli*-induced bacterial peritonitis model. Mice were intraperitoneally challenged with *E. coli* and intravenously treated with free Netil or 2-, 4-, or 8-arm Netil at 1 and 12 h after bacterial challenge, followed by survival monitoring for 168 h (**Figure 5a**). Untreated infected mice rapidly succumbed to bacterial peritonitis, whereas Netil-based treatments improved survival to different extents (**Figure 5b**). Among these groups, 8-arm Netil provided the strongest protection and achieved complete survival over the observation period, supporting its superior therapeutic activity in acute bacterial infection (**Figure 5b**). Survival differences were evaluated by Kaplan-Meier analysis with the log-rank test (n =10 mice per group; P <0.0001 for 8-arm Netil vs. untreated infected control). Consistently, body-weight monitoring showed improved recovery after 8-arm Netil treatment compared with untreated infected mice and lower-valency Netil conjugates **(Figure S15a-f)**.

We next examined whether this survival benefit was associated with reduced NET accumulation and inflammatory activation. *E. coli* challenge markedly increased NET levels in both blood and peritoneal fluid, indicating infection-induced NET formation and systemic dissemination of NET-associated inflammatory signals (**Figure 5c, d)**. Treatment with 8-arm Netil substantially reduced NET accumulation in both compartments, outperforming free Netil and lower-valency conjugates (**Figure 5c, d)**. In parallel, 8-arm Netil markedly suppressed inflammatory cytokine production, including serum TNF-α and IL-6, as well as serum MCP-1 and peritoneal TNF-α, IL-6, and MCP-1 (**Figure 5e, f** and **Figure S15g-j)**, demonstrating attenuation of both systemic and local inflammatory responses during bacterial peritonitis. Representative colony images and quantitative CFU analysis further showed that 8-arm Netil significantly decreased bacterial burden in both peritoneal fluid and blood, indicating effective control of local infection and systemic bacterial dissemination (**Figure 5g-i** and **Figure S15k**). Together, these results demonstrate that 8-arm Netil protects against *E. coli*-induced bacterial peritonitis by simultaneously reducing bacterial burden, NET accumulation, and infection-associated inflammatory activation.

**Figure 5.**
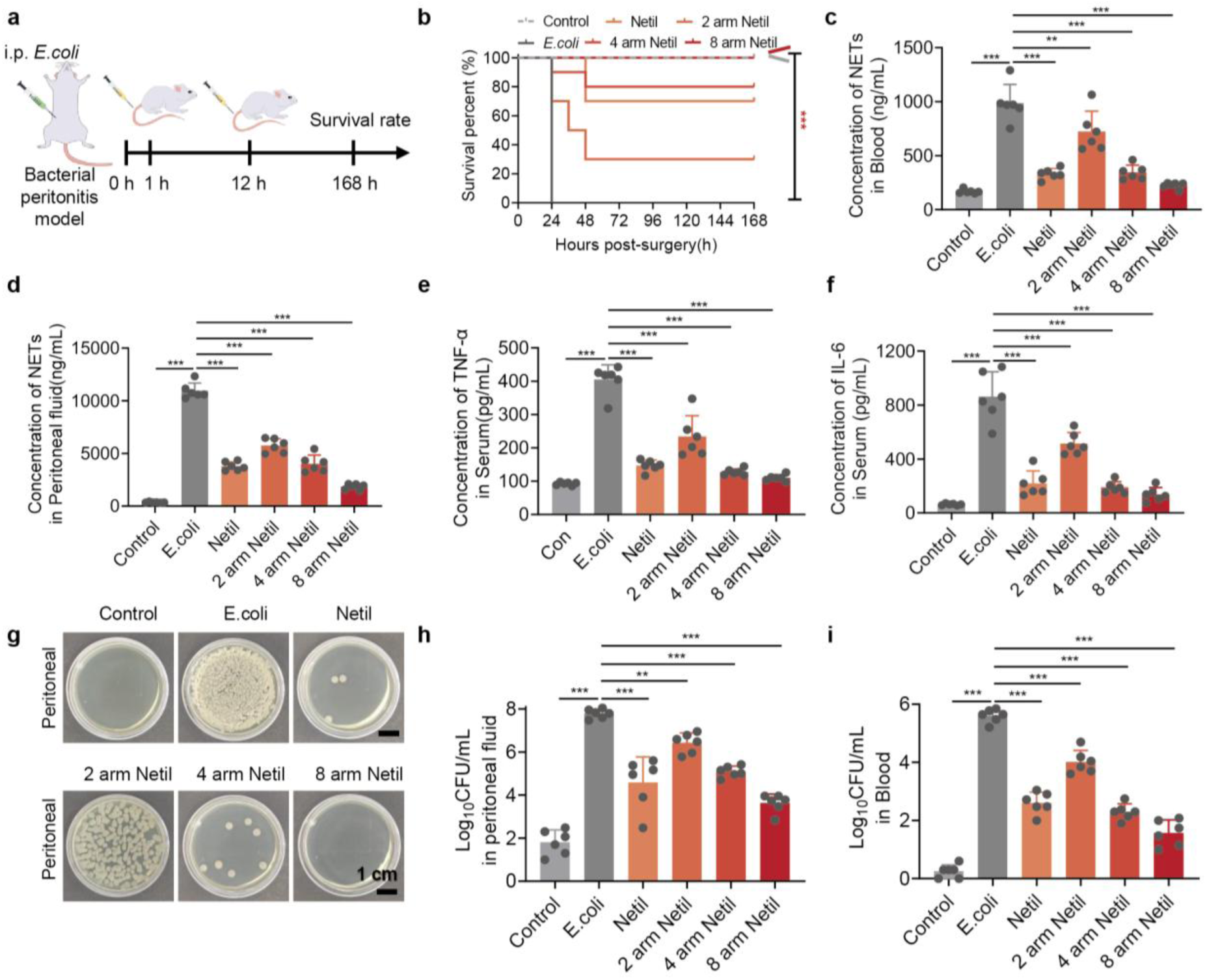
8-arm Netil protects against *E. coli*-induced bacterial peritonitis by reducing bacterial burden, NETs accumulation, and systemic inflammation. **(a)** Schematic illustration of the *E. coli*-induced bacterial peritonitis model and treatment schedule. Mice were challenged intraperitoneally with *E. coli* and treated intravenously with free Netil or 2-, 4-, or 8-arm Netil at 1 and 12 h after bacterial challenge. Survival was monitored for 168 h. **(b)** Survival of mice after *E. coli* challenge with different treatments. **(c, d)** Concentrations of NETs in blood **(c)** and peritoneal fluid **(d)** at 24 h after bacterial challenge. **(e, f)** Serum levels of TNF-α **(e)** and IL-6 **(f)** at 24 h after bacterial challenge. **(g)** Representative bacterial colony images from peritoneal fluid collected at 24 h after bacterial challenge. Scale bar, 1 cm. **(h, i)** Quantification of bacterial burden in peritoneal fluid **(h)** and blood **(i)** at 24 h after bacterial challenge. Data are presented as mean ± s.e.m. n = 6 biologically independent samples in **c-f, h,** and **i.** Statistical significance for quantitative data was calculated by one-way ANOVA with Tukey’s multiple-comparison test; survival was analyzed by Kaplan-Meier analysis with the log-rank test. **P < 0.01; ***P < 0.001. In **g,** representative images of bacteria colonies are shown (at least 6 images were taken for each group).

### 2.6 8-arm Netil protects against CLP-induced severe sepsis

To further evaluate the therapeutic efficacy of 8-arm Netil in polymicrobial severe sepsis, we used the cecal ligation and puncture (CLP) model, a clinically relevant model that recapitulates uncontrolled infection, systemic inflammation, and multiple-organ injury^[39]^. Mice received intravenous injections of DNase I, free Netil, or 2-, 4-, or 8-arm Netil at 1, 12, and 24 h after CLP, and survival was monitored for 168 h (**Figure 6a**). Untreated CLP mice showed rapid mortality and severe clinical deterioration, whereas 8-arm Netil provided the strongest survival protection among all treatment groups, achieving 100% survival over the observation period (**Figure 6b**). This survival benefit was accompanied by a marked reduction in clinical scores, indicating sustained improvement in disease severity after 8-arm Netil treatment (**Figure 6c**).

We next examined whether this protection was associated with reduced NET burden and inflammatory activation. CLP induced pronounced NET accumulation in both serum and peritoneal fluid, consistent with systemic and local release of NET-associated inflammatory signals during polymicrobial sepsis. DNase I partially reduced NET levels, particularly in the peritoneal compartment, but did not provide strong survival protection (**Figure 6b, d, e**). In contrast, 8-arm Netil substantially decreased NET levels in both serum and peritoneal fluid and showed stronger effects than free Netil, 2-arm Netil, or 4-arm Netil (**Figure 6d, e**). Immunofluorescence analysis further showed that 8-arm Netil accumulated in inflamed intestinal regions and reduced local NET deposition **(Figure S16)**, supporting its ability to access inflamed tissues and reduce both circulating and tissue-associated NET burden. Consistently, 8-arm Netil significantly reduced serum TNF-α, IL-6, IL-1β, and IL-17 compared with DNase I and free Netil (**Figure 6f-i**). These results suggest that DNA degradation alone or antibacterial activity alone is insufficient to fully suppress NET-associated inflammatory amplification, whereas 8-arm Netil attenuates systemic inflammation by coupling NET-associated chromatin scavenging with inflammatory signal suppression.

**Figure 6.**
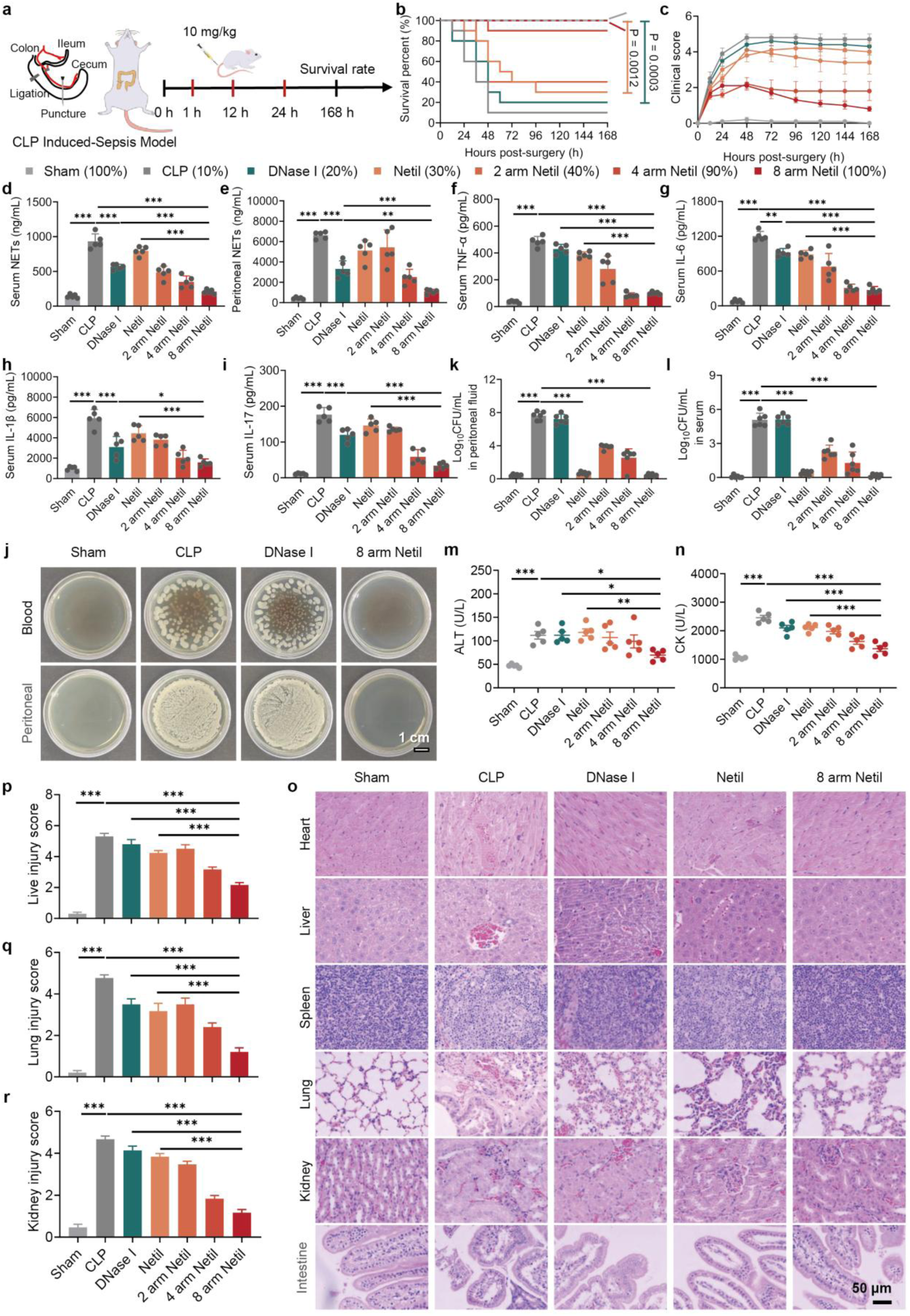
8-arm Netil protects against CLP-induced severe sepsis by reducing NETs, bacterial dissemination, systemic inflammation, and organ injury. **(a)** Schematic illustration of the cecal ligation and puncture (CLP)-induced sepsis model and treatment schedule. Mice received intravenous injections of DNase I, free Netil, or 2-, 4-, or 8-arm Netil at 1, 12, and 24 h after CLP, and survival was monitored for 168 h. **(b, c)** Survival curves **(b)** and clinical scores **(c)** of CLP-challenged mice after different treatments (n=6 independent mice). **(d, e)** NETs concentrations in serum **(d)** and peritoneal fluid **(e)** at 24 h after CLP. **(f-i)** Serum levels of TNF-α **(f)**, IL-6 **(g)**, IL-1β **(h)**, and IL-17 **(i)** at 24 h after CLP. **(j)** Representative bacterial colony images from blood and peritoneal fluid collected at 24 h after CLP. Scale bar, 1 cm. **(k, l)** Quantification of bacterial burden in peritoneal fluid **(k)** and serum **(l)** at 24 h after CLP. **(m, n)** Serum levels of alanine aminotransferase (ALT; **m)** and creatine kinase (CK; **n)** at 24 h after CLP. **(o)** Representative hematoxylin and eosin (H&E)-stained images of major organs collected at 24 h after CLP. Scale bar, 50 μm. **(p-r)** Quantitative injury scores of liver **(p)**, lung **(q)**, and kidney **(r)** at 24 h after CLP. Data are presented as mean ± s.e.m. n=5 independent mice in **d-l,** and **p-r**. Survival was analyzed by Kaplan-Meier analysis with the log-rank test. Statistical significance for other quantitative data was calculated by one-way ANOVA with Tukey’s multiple-comparison test; *P < 0.05, **P < 0.01, and ***P < 0.001. In **j,** representative images of bacteria colonies are shown (at least 6 images were taken for each group).

Because bacterial dissemination is a key driver of CLP-induced lethality^[40]^, we further quantified bacterial burden in blood and peritoneal fluid. Untreated and DNase I-treated CLP mice showed heavy bacterial growth in both compartments, whereas 8-arm Netil markedly reduced recoverable bacteria, indicating effective control of local polymicrobial infection and systemic bacterial dissemination (**Figure 6j-l** and **Figure S18)**. Consistent with reduced NET accumulation, cytokine production, and bacterial burden, 8-arm Netil also alleviated CLP-induced organ injury. Serum injury markers, including ALT, CK, AST, BUN, creatinine, and LDH, were reduced after 8-arm Netil treatment (**Figure 6m, n** and **Figure S19)**. Histological analysis showed that 8-arm Netil preserved tissue architecture and reduced inflammatory damage in the heart, liver, spleen, lung, kidney, and intestine (**Figure 6o-r** and **Figure S17)**.

Finally, long-term histological and serum biochemical analyses showed no obvious pathological abnormalities or sustained organ toxicity in surviving mice after 8-arm Netil treatment **(Figure S20)**. Together, these results demonstrate that 8-arm Netil provides robust protection against CLP-induced severe sepsis by integrating antibacterial activity, NET-associated chromatin scavenging, systemic cytokine suppression, and multi-organ protection while maintaining a favorable safety profile.

### 2.7 8-arm Netil suppresses inflammatory signaling in CLP-induced sepsis

Having demonstrated that 8-arm Netil reduced bacterial dissemination, NET accumulation, systemic cytokine production, and organ injury in CLP-induced severe sepsis, we next investigated whether these therapeutic effects were accompanied by restoration of immune homeostasis. Multiparametric flow cytometry was first used to profile circulating immune-cell populations at 24 h after CLP. t-distributed stochastic neighbor embedding (t-SNE) visualization showed pronounced remodeling of the peripheral immune landscape after CLP, with clear shifts in neutrophils, monocytes, CD4^+^ T cells, CD8^+^ T cells, NK cells, B cells, and other immune-cell populations compared with Sham mice (**Figure 7a**). Quantitative analysis further confirmed that CLP disrupted circulating immune-cell composition, whereas 8-arm Netil treatment partially restored the immune-cell distribution toward the Sham state (**Figure 7b** and **Figure S21)**. Consistent immune-regulatory effects were also observed in the peritoneal fluid, where 8-arm Netil modulated the proportions of neutrophils, macrophages, NK cells, B cells, total T cells, CD4^+^ T cells, and CD8^+^ T cells after CLP challenge **(Figure S22)**. These results indicate that 8-arm Netil does not merely suppress inflammation, but broadly rebalances immune-cell composition in both systemic and local inflammatory compartments.

To further decipher the potential mechanism by which the 8 arm Netil maintains immune homeostasis, we performed RNA-seq of circulating immune cells 24 h after CLP challenge. Venn analysis revealed overlapping and treatment-specific differentially expressed genes among sham, CLP, and 8-arm Netil-treated groups, indicating substantial transcriptional reprogramming after treatment (**Figure 7c** and **Figure S23)**. Direct comparison between CLP and 8-arm Netil-treated mice identified a large set of differentially expressed genes, including both downregulated and upregulated gene clusters (**Figure 7d**). KEGG enrichment analysis showed that TLR, MAPK, NF-κB, chemokine signaling pathways and the cytokine-cytokine receptor interaction both played important roles in the immunomodulation of the 8 arm Netil (**Figure 7e** and **Figure S23)**. Heatmap clustering further showed that 8-arm Netil reversed a broad CLP-induced inflammatory transcriptional signature, shifting gene-expression patterns closer to those observed in Sham mice (**Figure 7f** and **Figure S24)**. Similar pathway-level regulation was also observed in peritoneal fluid transcriptomic analyses, supporting coordinated systemic and local immune modulation **(Figure S25)**.

**Figure 7.**
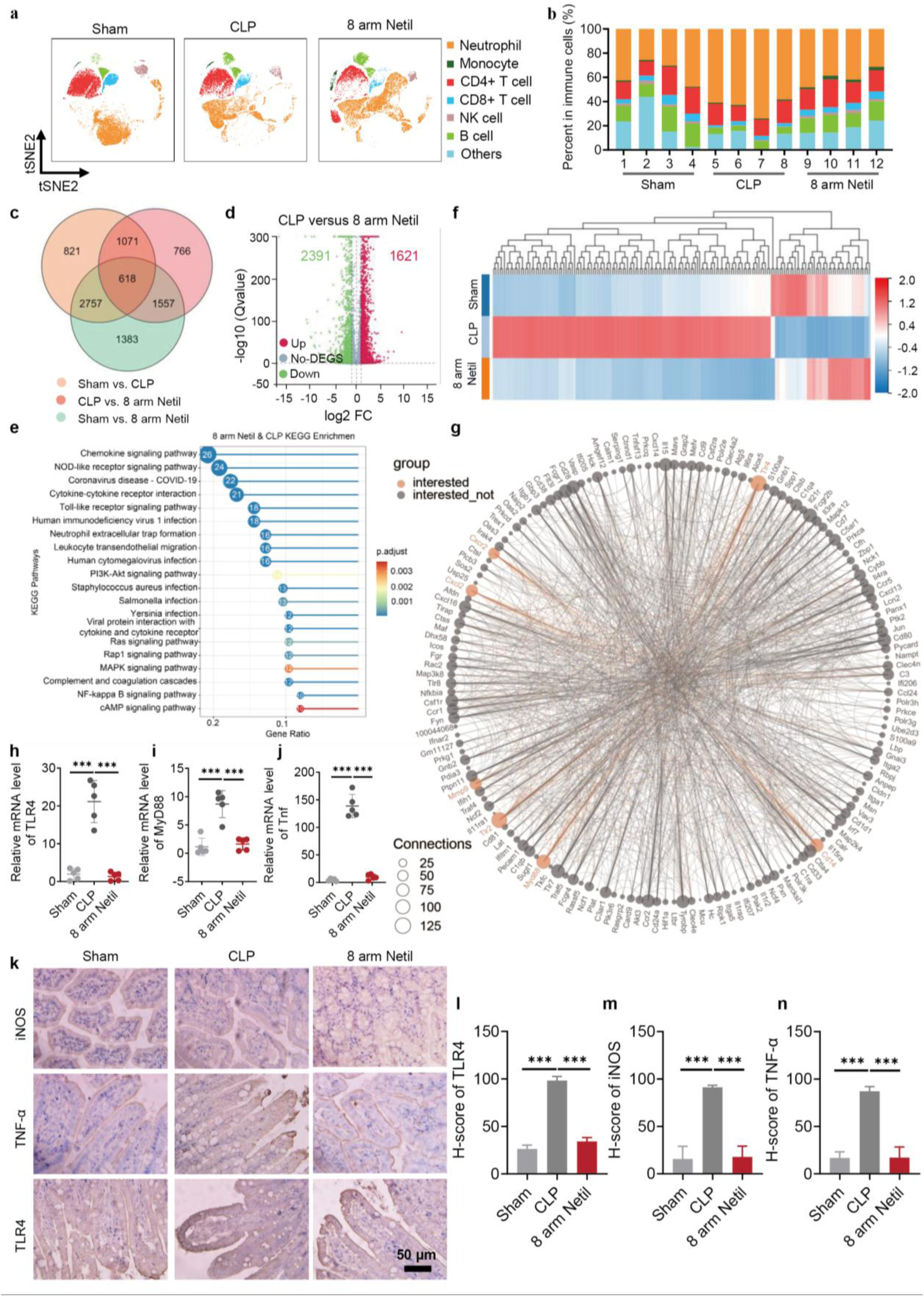
8-arm Netil suppresses inflammatory signaling in CLP-induced sepsis. **(a)** t-SNE visualization of circulating immune cell populations analyzed by multiparametric flow cytometry at 24 h after CLP. Major immune cell subsets, including neutrophils, monocytes, CD4^+^ T cells, CD8^+^ T cells, NK cells, B cells, and other immune cells, are indicated by distinct colors. **(b)** Quantification of immune cell subset proportions in blood from Sham, CLP, and 8-arm Netil-treated mice at 24 h after CLP. **(c)** Venn diagram showing overlapping and distinct differentially expressed genes (DEGs) among Sham, CLP, and 8-arm Netil-treated groups. **(d)** Volcano plot of DEGs between CLP and 8-arm Netil-treated mice. Upregulated and downregulated genes are shown in red and green, respectively. **(e)** KEGG pathway enrichment analysis of DEGs between CLP and 8-arm Netil-treated mice, highlighting inflammatory and immune-related pathways regulated by 8-arm Netil. **(f)** Heatmap showing hierarchical clustering of representative DEGs in sham, CLP, and 8-arm Netil-treated groups. **(g)** Protein-protein interaction network of DEGs between CLP and 8-arm Netil-treated groups. Highlighted nodes indicate selected inflammation-related genes, and node size reflects the number of protein connections. **(h-j)** qRT-PCR validation of representative inflammatory genes, including TLR4 **(h)**, MyD88 **(i)**, and Tnf **(j)**, in blood at 24 h after CLP. **(k)** Representative immunohistochemistry images of iNOS, TNF-α, and TLR4 in colon tissues collected at 24 h after CLP. Scale bar, 50 μm. **(l-n)** Quantification of immunohistochemistry staining for TLR4 **(l)**, iNOS **(m)**, and TNF-α **(n)** in colon tissues. In **b**, the data are presented as the mean (n=4 independent mice per group). In **h-j** and **l-n**, data are presented as mean ± s.e.m. (n=5 biologically independent samples in **h-j;** n=4 biologically independent samples in **l-n)**. Statistical significance was calculated by one-way ANOVA with Tukey’s multiple-comparison test; ***P < 0.001.

Protein-protein interaction analysis of differentially expressed genes (DEGs) further highlighted inflammation-related molecular networks affected by 8-arm Netil treatment (**Figure 7g**). Several highly connected nodes were associated with innate immune activation, cytokine signaling, chemokine responses, Toll-like receptor signaling, and NET-associated inflammatory amplification, suggesting that 8-arm Netil attenuates interconnected inflammatory modules rather than a single downstream cytokine. qRT-PCR validation confirmed that CLP strongly increased the expression of representative inflammatory genes, including *Tlr4*, *Myd88*, and *Tnf*, whereas 8-arm Netil markedly suppressed their expression in blood (**Figure 7h-j** and **Figure S26)**. These transcriptional data are consistent with the earlier observation that 8-arm Netil neutralizes NET-induced TLR4/TLR9 activation *in vitro* and reduces systemic cytokine levels *in vivo* (**Figure 4h-p** and **Figure 6f-i**). Additionally, 8 arm Netil sharply decreased the colonic expression of iNOS, TNF-α, and TLR4 (**Figure 7k-n** and **Figure S27).** Together, these results demonstrate that 8-arm Netil suppresses inflammatory signaling at both transcriptional and tissue levels.

### 2.8 8-arm Netil shows prolonged circulation, inflammation-associated tissue accumulation, and favorable biosafety

Finally, we evaluated the pharmacokinetic behavior and tissue distribution of 8-arm Netil after intravenous administration. 8-arm Netil showed relatively slow blood clearance, with 93.4% and 18.3% of the injected dose remaining in circulation at 1 and 24 h, respectively (**Figure 8a**). Its elimination half-life (T_1/2_ = 4.0 h) was 2.67-fold longer than that of free Netil (T_1/2_ = 1.5 h), consistent with the prolonged circulation expected from its macromolecular architecture (**Figure 8a**). This extended blood residence may provide a broader therapeutic window for capturing circulating NET-associated DNA-histone chromatin complexes and accessing inflamed tissues during sepsis. Biodistribution analysis further showed that 8-arm Netil had higher blood retention and greater accumulation in inflamed intestinal tissues than free Netil and lower-valency Netil conjugates, while maintaining relatively limited kidney accumulation (**Figure 8b**). This profile is favorable because the intestine is a major inflammatory and barrier-injury site in CLP-induced sepsis, whereas renal accumulation is a key concern for aminoglycoside-associated toxicity. Consistently, ICG-labeled 8-arm Netil showed stronger intestinal fluorescence in CLP mice than in healthy controls over 2-24 h, supporting inflammation-associated tissue accumulation after systemic administration (**Figure 8c-h**).

We next assessed whether the prolonged circulation and tissue accumulation of 8-arm Netil caused detectable systemic toxicity. Hematological analysis showed no obvious changes in white blood cells, red blood cells, neutrophils, lymphocytes, monocytes, or platelets after 8-arm Netil treatment in healthy mice (**Figure 8i**). Consistently, hemocompatibility assays showed minimal hemolysis and preserved red blood cell morphology across the tested concentration range **(Figure S28)**. Serum biochemical markers, including LDH, CK, BUN, ALT, AST, and creatinine, remained comparable between control and 8-arm Netil-treated mice, indicating no evident cardiac, hepatic, or renal toxicity under the tested conditions (**Figure 8j**). Histological examination further showed preserved tissue architecture in the heart, liver, spleen, lung, kidney, and cecum after 8-arm Netil administration (**Figure 8k**). Together, these hematological, hemocompatibility, biochemical, and histological analyses support the favorable systemic biosafety of 8-arm Netil after intravenous administration.

**Figure 8.**
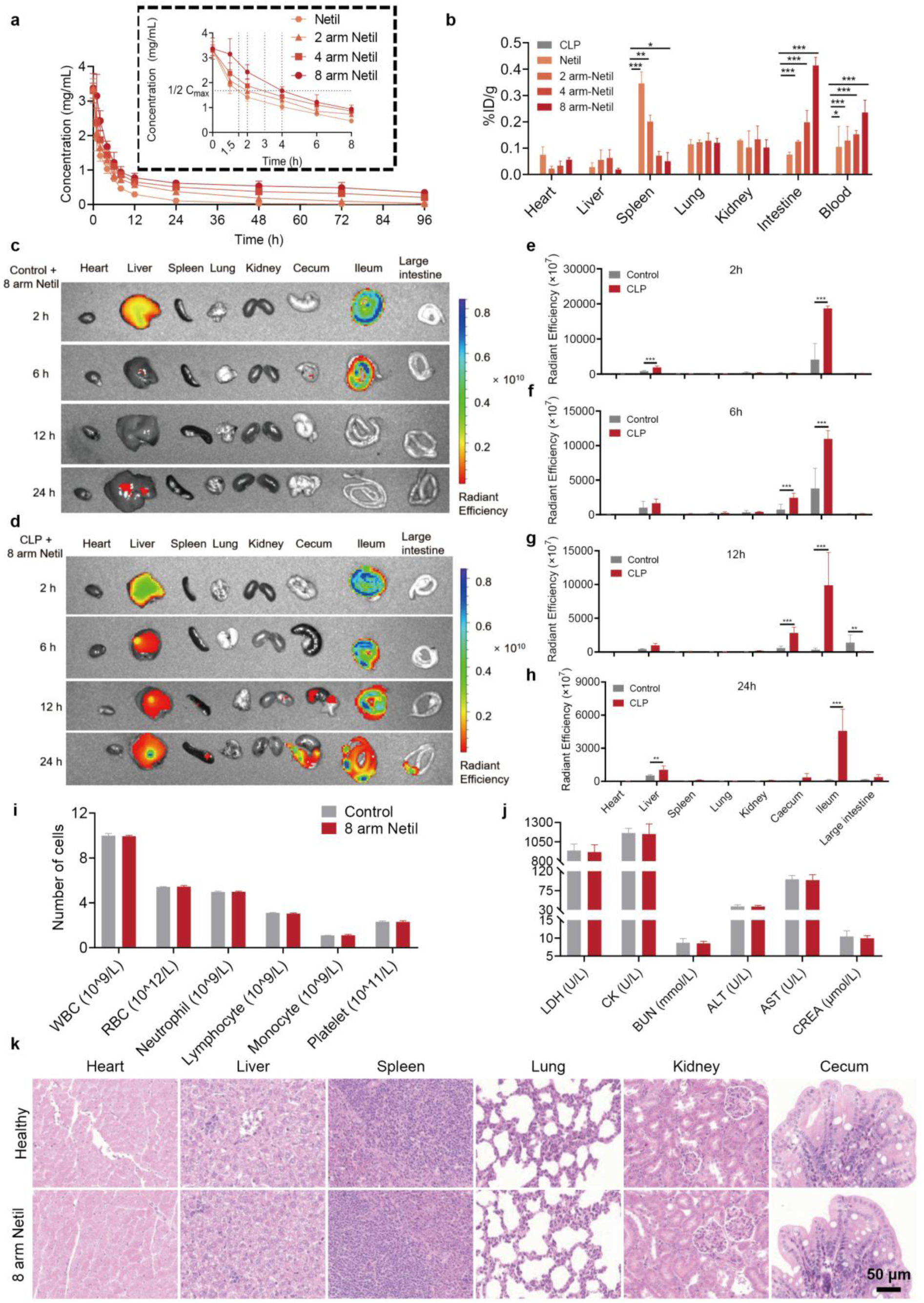
Pharmacokinetics, biodistribution, and biosafety evaluation of 8-arm Netil. **(a)** Pharmacokinetic profiles of free Netil and 2-, 4-, and 8-arm Netil after intravenous administration in BALB/c mice. The inset shows the early concentration-time profiles. **(b)** Quantitative biodistribution of Netil and 2-,4-, and 8-arm Netil in major organs and blood at 24 h after intravenous administration, as determined by inductively coupled plasma optical emission spectrometry (ICP-OES) using Ru labeling and expressed as percentage of injected dose per gram tissue (%ID/g). **(c, d)** Ex vivo fluorescence imaging of major organs and intestinal tissues from control mice **(c)** and CLP-induced septic mice **(d)** at 2, 6, 12, and 24 h after intravenous administration of ICG-labeled 8-arm Netil. **(e-h)** Quantification of fluorescence signals in major organs and intestinal tissues at 2 h **(e)**, 6 h **(f)**, 12 h **(g)**, and 24 h **(h)** after administration of labeled 8-arm Netil in control and CLP-induced septic mice. **(i)** Hematological analysis of white blood cells, red blood cells, neutrophils, lymphocytes, monocytes, and platelets in healthy mice after 8-arm Netil treatment. **(j)** Serum biochemical analysis of lactate dehydrogenase (LDH), creatine kinase (CK), blood urea nitrogen (BUN), alanine aminotransferase (ALT), aspartate aminotransferase (AST), and creatinine (CREA) in healthy mice after 8-arm Netil treatment. **(k)** Representative H&E-stained images of major organs and cecum from healthy control mice and 8-arm Netil-treated mice. Scale bar, 50 μm. Data are presented as mean ± s.e.m. (n = 3 independent mice or experiments). Statistical significance was calculated by one-way ANOVA with Tukey’s multiple-comparison test; *P < 0.05, **P < 0.01, and ***P < 0.001.

## 3. Conclusion

In summary, we developed a multivalent macromolecular aminoglycoside platform that integrates antibacterial activity with NET-associated chromatin scavenging for severe sepsis treatment. By conjugating aminoglycosides onto 2-, 4-, and 8-arm PEG, we established a valency-tunable NET-neutralizing material library and identified 8-arm Netil as the lead construct. Compared with free aminoglycosides, 8-arm Netil retained antibacterial function while gaining high-avidity and selective binding to NET-associated DNA-histone chromatin complexes, thereby addressing two coupled pathological drivers of severe sepsis: uncontrolled infection and persistent inflammatory signal amplification.

A central material-driven advance of this work is the transformation of a conventional antibacterial molecule into a chromatin-targeting NET neutralizer through multivalent scaffold engineering. The 8-arm architecture enabled stable multivalent noncovalent interactions with intact DNA-histone chromatin complexes and showed stronger recognition of assembled chromatin than isolated DNA, histones, or abundant serum proteins. This binding mode allowed 8-arm Netil to directly complex with reconstituted chromatin and native NETs, thereby suppressing NET-induced TLR4/TLR9 activation and macrophage inflammatory responses. These findings establish NET-associated DNA-histone chromatin complexes as actionable extracellular targets and highlight multivalent chromatin recognition as a rational design principle for immunomodulatory biomaterials.

In both *Escherichia coli*-induced bacterial peritonitis and CLP-induced severe sepsis, intravenously administered 8-arm Netil reduced bacterial dissemination, NET accumulation, systemic cytokine production, and multiple-organ injury, leading to markedly improved survival under the treatment schedules and challenge severities tested in this study. Beyond acute inflammatory suppression, 8-arm Netil also rebalanced immune-cell composition and attenuated inflammatory signaling in systemic and tissue compartments, supporting immune-homeostatic restoration rather than simple cytokine inhibition. Its prolonged circulation, inflammation-associated tissue accumulation, hemocompatibility, and favorable hematological, biochemical, and histological safety profiles further support its suitability for systemic administration in severe infection-associated inflammation.

Overall, this study provides a materials-based strategy for converting antibacterial small molecules into multifunctional macromolecular therapeutics capable of coordinating pathogen control with inflammation resolution. Beyond severe sepsis, multivalent DNA-histone chromatin complex scavenging may be broadly applicable to other NET-driven infectious and inflammatory diseases.

## 4. Experimental Section/Methods

### Synthesis of macromolecular aminoglycosides

Macromolecular aminoglycosides were prepared through a two-step conjugation strategy. First, 8-arm PEG-OH was carboxylated with suberic anhydride to obtain 8-arm PEG-COOH. The resulting carboxylated PEG was then coupled with aminoglycosides to afford 8-arm aminoglycoside conjugates (8-arm AG). The synthesis of 8-arm Netil is described as a representative example. Suberic anhydride was prepared and purified according to a reported procedure^[41]^. 8-arm PEG-OH (1 g, 0.1 mmol) and suberic anhydride (624 mg, 4 mmol) were dissolved in anhydrous N, N-dimethylformamide (DMF, 5 mL), followed by the addition of triethylamine (224 μL, 1.6 mmol). The reaction mixture was stirred at room temperature for 24 h, quenched with 5% hydrochloric acid aqueous solution, and extracted with dichloromethane (DCM). The combined organic phase was concentrated under reduced pressure and purified by silica gel flash chromatography to yield 8-arm PEG-COOH as a yellow oil. For Netil conjugation, netilmicin (238 mg, 0.5 mmol) and triethylamine (14 μL, 0.1 mmol) were dissolved in dimethyl sulfoxide (DMSO, 8 mL) and stirred at room temperature. A solution of 8-arm PEG-COOH (50 mg, 0.005 mmol) in DMSO (2 mL) was added dropwise, and the reaction was continued for 24 h at room temperature. The crude product was concentrated and purified by flash chromatography to afford 8-arm Netil as a yellow solid. The average number of free versus conjugated netilmicin primary amines was determined by ¹H NMR integration and a quantitative ninhydrin assay against a netilmicin standard curve, to confirm that the cationic groups required for antibacterial activity were not fully consumed during amide coupling. The number-average molecular weight (Mn) and weight-average molecular weight (Mw) were determined by gel permeation chromatography (GPC). The chemical structure was characterized by ¹H NMR spectroscopy using D₂O as the solvent.

### Cells

Mouse neutrophils were isolated by Percoll density-gradient centrifugation and maintained in DMEM supplemented with 10% fetal bovine serum (FBS) at 37 °C in 5% CO₂. Human umbilical vein endothelial cells (HUVECs) and RAW264.7 macrophages were cultured under the same conditions. RAW264.7 cells between passages 3 and 7 were used. HEK-Blue TLR4 and TLR9 reporter cells were cultured in DMEM containing 10% FBS and 1% penicillin-streptomycin at 37 °C in 5% CO₂ and used between passages 2 and 10.

### Patient samples

We recruited 40 healthy volunteers and 100 sepsis patients from the ICUs at the First Affiliated Hospital of Zhengzhou University. Participants were at least 18 years old, with consent obtained directly or via a surrogate. Sepsis patients met the Sepsis-3 criteria^[3]^, had a SOFA score ≥2 following a bacterial infection. Consent and sample collection occurred within 72 hours of ICU admission. Whole blood was collected, centrifuged, and stored at -80°C for future analysis. Healthy volunteers had no infections or immune disorders and were at least 18 years old. This study was approved by the hospital’s Ethics Committee (approval no. KY-2022-2768).

### Animals

All animal experiments were performed according to protocols approved by the Laboratory Animal Ethics Committee of South China University of Technology (Guangzhou, China). Male BALB/c mice aged 4-6 weeks were purchased from Hunan SJA Laboratory Animal Co., Ltd. All animals were acclimatized for at least 1 week before experiments.

### Cytocompatibility and hemocompatibility

The cytocompatibility of free aminoglycosides and 8-arm AG was evaluated in RAW264.7 cells and HUVECs using the MTT assay after material incubation. Cell viability was calculated from absorbance at 570 nm. In vitro hemocompatibility was assessed by incubating red blood cells with free aminoglycosides, 8-arm aminoglycoside conjugates, or PEI25k at different concentrations. PBS and deionized water served as negative and positive controls, respectively. Hemolysis was quantified from absorbance at 545 nm, and red blood cell morphology was examined by scanning electron microscopy.

### In vitro antibacterial activity

Minimum inhibitory concentration (MIC) and minimum bactericidal concentration (MBC) assays were performed to evaluate the antibacterial activity of free aminoglycosides and 8-arm aminoglycoside conjugates against *Escherichia coli* (ATCC 25922). Bacteria were incubated with serially diluted materials in LB medium, and bacterial growth was assessed by OD600 as reported method^[42]^. For bactericidal analysis, aliquots from the MIC assay were plated on LB agar, and colony formation was examined after incubation.

### Preparation and characterization of extracellular chromatin

Extracellular chromatin was prepared by assembling histones with DNA, including enriched CpG DNA, at different concentrations and molar ratios. The optimized formulation consisted of histone (30 μg/mL) and enriched CpG DNA (10 μg/mL) at an equal molar ratio, which induced robust activation of immune reporter cells and macrophages. DNase I (5 U/mL) was used to confirm the DNA-dependent immunostimulatory activity of extracellular chromatin. Morphology was examined by scanning electron microscopy.

### Neutrophil isolation and n-NET purification

Neutrophils were isolated from mouse bone marrow or rat peripheral blood using Percoll density-gradient centrifugation^[43]^. Isolated neutrophils were resuspended in DMEM supplemented with 10% fetal bovine serum and maintained at 37 °C in 5% CO2. To generate native NETs (n-NETs), neutrophils were stimulated with phorbol 12-myristate 13-acetate (PMA, 100 nM) for 4 h at 37 °C. NET-containing fractions were collected, cleared of cellular debris, concentrated by centrifugation, and resuspended in PBS. NET concentration was determined based on DNA content using the Quant-iT PicoGreen dsDNA assay and fluorescence detection at Ex/Em = 480/520 nm, as previously described^[44]^. Purified n-NETs were stored at -80 °C until use.

### Scanning electron microscopy

Purified n-NETs or extracellular chromatin samples were fixed with 2.5% glutaraldehyde, washed with PBS, dehydrated through a graded ethanol series, dried on support films, and sputter-coated with platinum. Sample morphology was examined by scanning electron microscopy using a Q25 microscope.

### In vitro scavenging of danger signals

Direct binding of danger signals was evaluated according to a reported method with minor modifications^[45]^. Briefly, extracellular chromatin, LPS, HMGB1, or TNF-α was incubated with 8-arm aminoglycoside conjugates at room temperature for 3 h. For extracellular chromatin binding, chromatin was fixed at 20 μg/mL and mixed with materials at chromatin-to-material mass ratios of 1:0.5, 1:1, 1:2, 1:4, 1:8, 1:16, 1:32, and 1:64, corresponding to material concentrations of 10-1280 μg/mL. For other danger signals, LPS (200 ng/mL), HMGB1 (20 μg/mL), or TNF-α (20 ng/mL) was incubated with 8-arm aminoglycoside conjugates (200 μg/mL). After centrifugation, residual danger signals in the supernatant were quantified using the corresponding assays. Extracellular chromatin was measured using the Quant-iT PicoGreen dsDNA assay, LPS using a Limulus amebocyte lysate endotoxin quantification kit, and HMGB1 and TNF-α using ELISA kits. Binding efficiency was calculated from the change in danger-signal concentration before and after material incubation.

### Serum danger-signal scavenging

Serum samples from CLP-induced septic mice or patients with sepsis were incubated with 8-arm aminoglycoside conjugates (200 μg/mL) at room temperature for 3 h. After centrifugation, residual n-NETs, LPS, HMGB1, and TNF-α in the supernatant were quantified using the assays described above.

### In vitro anti-inflammatory assays

HEK-Blue TLR4/TLR9 reporter cells or RAW264.7 macrophages were seeded in 96-well plates and stimulated with sepsis-related inflammatory stimuli, including serum from CLP-induced septic mice or patients with sepsis, n-NETs, extracellular chromatin, or 8-arm Netil was then added at the indicated concentrations. After 24 h, TLR4/TLR9 reporter activation was quantified using QUANTI-Blue by measuring secreted embryonic alkaline phosphatase activity at 620 nm. Macrophage activation was assessed by measuring TNF-α levels in the culture supernatant using ELISA.

### Immunofluorescence staining and colocalization

Purified n-NETs were stained with antibodies against citrullinated histone H3 and myeloperoxidase, followed by fluorescent secondary antibodies and DAPI counterstaining. Fluorescent extracellular chromatin was prepared by assembling DAPI-labeled DNA with Cy5-labeled histone. FITC-labeled 8-arm Netil was incubated with n-NETs or extracellular chromatin, and the resulting complexes were collected for fluorescence imaging. For cellular colocalization, labeled n-NETs and FITC-labeled 8-arm Netil were incubated with RAW264.7 macrophages, followed by lysosome (LysoTracker Red DND-99) and nuclear (DAPI) staining. Images were acquired by fluorescence microscopy or confocal laser scanning microscopy.

### ITC assay

Binding thermodynamics between materials and biological substrates were measured using a MicroCal PEAQ-ITC system at 25 °C. PAMAM G4, Netil, 2-arm Netil, 4-arm Netil, and 8-arm Netil were tested against extracellular chromatin, histone, DNA, or serum albumin (SA). For a representative assay, 8-arm Netil solution (7.24 × 10^-5^ mol/L, 80 μL) was loaded into the injection syringe and titrated into substrate solution (1.5 × 10^-5^ mol/L, 300 μL) in the sample cell. Titrations were performed with 19 injections of 2.0 μL at a stirring speed of 750 rpm. Background heat from blank titrations was subtracted before fitting the binding parameters.

### Molecular modeling and dynamics simulation

The extracellular chromatin model was constructed from histone and DNA fragments based on the experimental DNA sequence and known histone-DNA structural information (PDB ID: 6PWW). Docking was performed using HADDOCK and molecular interaction sites were further evaluated using Molecular Operating Environment. Molecular dynamics simulations were conducted using GROMACS with CHARMM-based force fields. Netil and 8-arm Netil structures were generated, energy-minimized, docked to the chromatin model, and simulated to evaluate binding stability. Interaction energies were estimated by the molecular mechanics/Poisson-Boltzmann surface area (MM/PBSA) method over the equilibrated portion of each trajectory. Simulation results were visualized using PyMOL.

### Bacterial peritonitis model

*E. coli* was cultured overnight in LB medium at 37 °C, collected by centrifugation, washed with saline, and resuspended to 1.5 × 10^9^ CFU/mL. Male BALB/c mice were intraperitoneally challenged with 200 μL of the bacterial suspension. At 1 and 12 h after infection, mice received intravenous injections of saline, free Netil (2.75 mg/kg⁻¹), or 2-, 4-, 8-arm Netil (10 mg/kg). Survival, body weight, bacterial burden in blood and peritoneal fluid, NET levels, and inflammatory cytokines were analyzed at the indicated time points.

### CLP-induced sepsis models and treatment

Cecal ligation and puncture was performed as previously described^[10]^. The cecum was ligated and punctured twice with a 21-gauge needle. Sham mice underwent the same procedure without ligation or puncture. Mice were randomized into treatment groups and intravenously administered saline, free Netil, 8-arm Netil, DNase I, or other comparator treatments at the indicated doses and time points. Survival and clinical scores were monitored for 168 h.

### Survival rate and Clinical score

Mice were monitored for survival rate and clinical score for 168h. For the clinical score, mice were scored every 12 h using the criteria as previously described1: score 0, no symptoms; score 1, piloerection and huddling; score 2, piloerection, diarrhea, and huddling; score 3, lack of interest in surroundings and severe diarrhea; score 4, decreased movement and listless appearance; and score 5, loss of self-righting reflex. Mice were humanely euthanized when they exhibited a score of 5.

### In vivo bacterial burden

At the indicated time points after CLP or bacterial peritonitis challenge, blood and peritoneal fluid were collected. Samples were serially diluted, plated on LB agar, and incubated to quantify colony-forming units.

### Serum biochemical and cytokine analysis

Blood was collected after CLP or bacterial challenge, and serum was isolated for biochemical and cytokine analysis. Serum ALT, AST, creatinine, urea, BUN, LDH, and CK were measured using an automatic biochemical analyzer. Cytokines, including TNF-α, IL-6, MCP-1, IL-1β, and IL-17, were quantified using ELISA kits according to the manufacturers’ instructions as reported method^[46]^.

### Histology and immunohistochemistry

Major organs were collected, fixed, embedded, sectioned, and stained with hematoxylin and eosin. Histological injury was evaluated in a blinded manner according to reported criteria^[10]^. For immunohistochemistry, colon sections were stained with antibodies against TLR9, TLR4, TNF-α, iNOS, CD206, and TGF-β, followed by DAB development and hematoxylin counterstaining. Positive staining was quantified using H-score analysis and ImageJ. For tissue immunofluorescence, frozen liver and colon sections were stained with antibodies against citrullinated histone H3, or F4/80, followed by fluorescent secondary antibodies and DAPI counterstaining.

### Flow cytometric analysis

24 h post-CLP, mice in all groups were anesthetized. Whole blood was collected into pediatric heparin tubes and then centrifuged at 400 g for 15 min at 4°C to obtain the leukocyte. Afterward, mice were killed and then peritoneal fluid were harvested for obtained peritoneal cells. All obtained cells were resuspended with PBS after lysed by red blood cell lysis buffer (C3702, Beyotime Biotechnology) and then counted. Then cells were blocked with mice TruStain FcX^TM^ (anti-mouse CD16/32) for 10 min, 4°C before surface staining. Then cells were stained using a panel of antibodies as follows: FITC anti-mouse NK-1.1 Antibody (clone: PK136; Biolegend), PE/Cyanine7 anti-mouse F4/80 Antibody (clone BM8; Biolegend), PE/Cyanine5 anti-mouse CD19 Antibody (clone:6D5; Biolegend), Alexa Fluor® 647 anti-mouse/human CD44 Antibody (clone: IM7; Biolegend), APC/Cyanine7 anti-mouse CD45.2 Antibody (clone:104; Biolegend), Alexa Fluor® 700 anti-mouse I-A/I-E Antibody (Clone: M5/114.15.2; Biolegend), Brilliant Violet 421™ anti-mouse CD3 Antibody (clone: 17A2; Biolegend), Brilliant Violet 711™ anti-mouse CD8a Antibody (clone:53-6.7; Biolegend), PerCP/Cyanine5.5 anti-mouse CD11c Antibody (clone: N418; Biolegend), PE-CF594 Rat Anti-Mouse Ly-6G and Ly-6C Antibody (clone: RB6-8C5; BD), V500 Rat anti-CD11b Antibody (clone: M1/70; BD) and BUV563 Rat Anti-Mouse CD4 (clone: GK1.5; BD) for 30 min at 4°C. After antibody staining, cells were washed with PBS and centrifugated for 5 min at 400 g, then the supernatants were discarded and cells were resuspended with PBS. All samples were analyzed on a Fortessa (BD Biosciences) flow cytometer. Data analysis and graphical interpretation were performed with FlowJo software. T-distributed stochastic neighbor embedding (tSNE) was used for the analysis of the populations of distinct immune cells in blood and peritoneal fluid. viSNE analysis was run on 40000 live CD45.2+ single cells per sample using all markers.

### RNA-sequencing and data analysis

At 24 h post-CLP, mice in Sham, CLP, and 8 arm Netil groups were anesthetized. Whole blood was collected into pediatric heparin tubes and then mice were killed, and peritoneal fluid was harvested for obtained peritoneal cells. All obtained cells were resuspended by PBS after lysed by red blood cell lysis buffer and centrifugated for 5 min at 400 g, then the supernatants were discarded. After that, the cells were frozen with liquid nitrogen and transferred to -80 °C. All samples were sent to BGI for transcriptome sequencing. RNA quality was determined using an Agilent 2100 Bioanalyzer. Sequencing libraries were prepared using the Illumina TruseqTM RNA Library Prep Kit v2. For bioinformatics analysis, the expression level of each transcript was calculated according to the fragments per kilobase of exon per million mapped reads (FPKM) method. Differential expression analysis was performed using the DEGseq with |log2 FC| ≥ 0.5 and Q value ≤ 0.001, and Kegg enrichment analysis of annotated differentially expressed genes was performed by Phyper based on the Hypergeometric test. The significant levels of terms and pathways were corrected by Q value with a rigorous threshold by Bonferroni. Upon the selected DEGseq protein list, we extracted potential protein-protein interactions from the STRING database with the R package STRINGdb (v.2.4.2)46. 164 out of 171 genes were extracted in the STRING database by setting the score threshold of 400. The protein-protein interaction for all types between these 164 genes was extracted using the “map” method in STRINGdb. A total of 2510 interactions were mapped. The protein-protein interaction network was constructed with ggraph (v.2.0.5) and visualized with igraph (v.1.2.6).

### Quantitative real-time PCR analysis

At 24 h post-CLP, mice in sham, CLP, and 8 arm Netil groups were anesthetized. Whole blood was collected into pediatric heparin tubes and then mice were killed and peritoneal fluid were harvested for obtained peritoneal cells. All obtained cells were resuspended by PBS after lysed by red blood cell lysis buffer and centrifugated for 5 min at 400 g, then the supernatants were discarded. After that, the total RNA of blood or peritoneal fluid cells was extracted by TRIzol reagent (Invitrogen) as the manufacturer’s protocol. Next, reverse transcription of 1 μg RNA into cDNA was performed using the PrimeScriptTM RT reagent Kit with gDNA Eraser (Takara). The cDNA was amplified by PCR with TB Green® Premix Ex TaqTM Ⅱ Kit (Takara) in a real-time PCR system (LightCycler® 96 instrument, Roche) to determine the mRNA levels of TLR2, TLR4, MyD88, Mmp9, CD14, CXCL2, CXCR2, and Tnf. The amplified transcripts were quantified by the comparative 2-ΔΔCt method. The following primer sequences synthesized by Sangon Biotech were used: TLR2 forward:

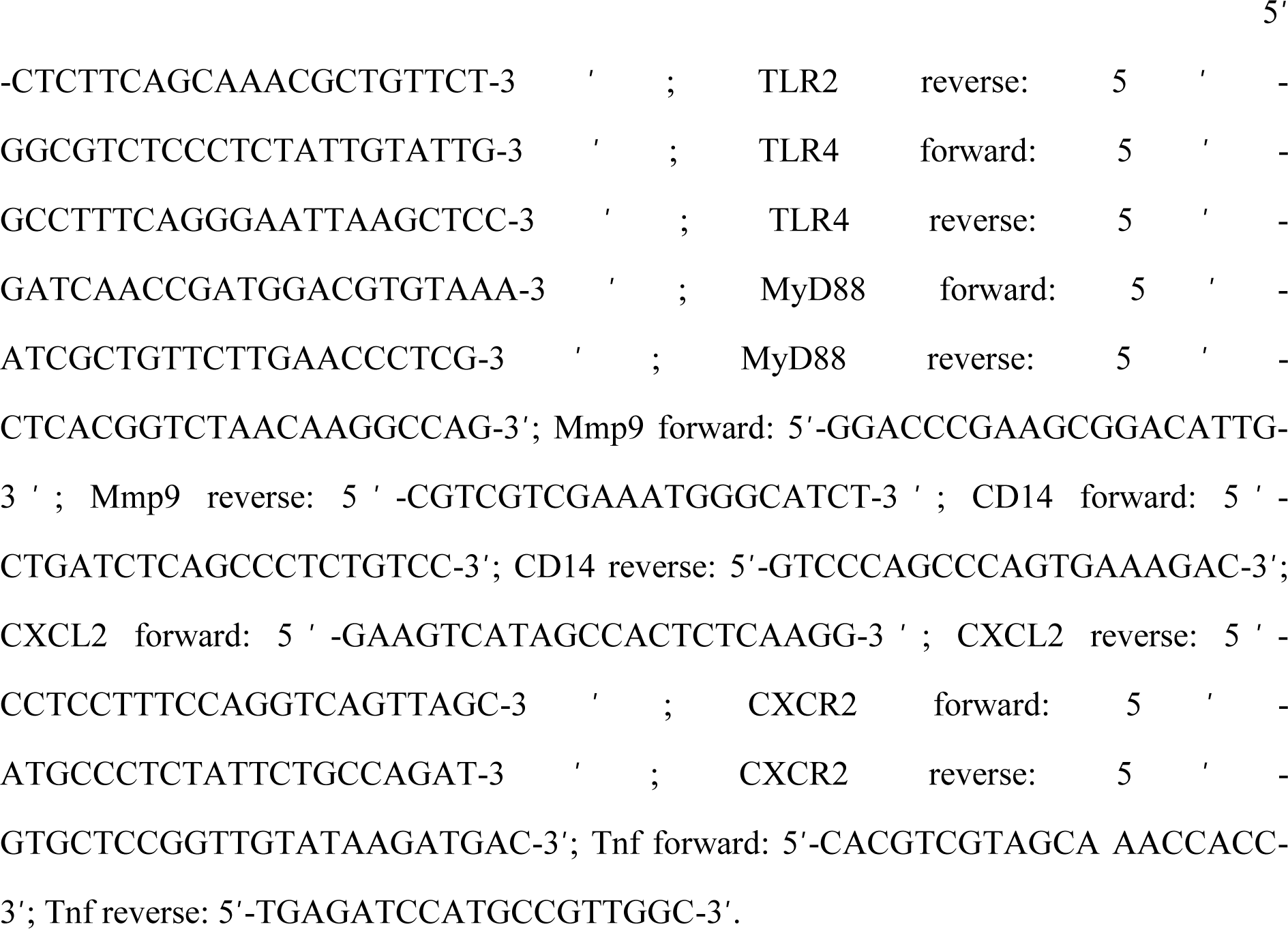

### Pharmacokinetics and tissue distribution

Ruthenium-labeled 8-arm Netil was used to evaluate the pharmacokinetics and tissue distribution of 8-arm Netil. Briefly, ruthenium(III) chloride (200 mg, 0.97 mmol) and 2,2′-bipyridine-4,4′-dicarboxylic acid (500 mg, 2.05 mmol) were dissolved in anhydrous DMF under an argon atmosphere and reacted at 170 °C for 3 h. The product was precipitated with acetone at -20 °C overnight, collected by filtration, and dried to obtain Ru(H_4_dcbpy)_4_Cl_2_. The ruthenium reagent was then prepared in deionized water and incubated with 8-arm Netil at 25 °C for 24 h to generate ruthenium-labeled 8-arm Netil. For pharmacokinetic analysis, ruthenium-labeled 8-arm Netil was intravenously injected into BALB/c mice, and blood samples were collected at predetermined time points after injection. For tissue distribution analysis, mice were sacrificed 24 h after administration, and blood and major organs, including the heart, liver, spleen, lung, kidney, and intestine, were collected and weighed. Samples were digested, and ruthenium content was quantified by inductively coupled plasma optical emission spectroscopy (ICP-OES).

### In vivo biosafety assessment

Long-term biosafety after sepsis treatment was evaluated in surviving mice from the CLP model. Sham mice and surviving mice treated with 4-arm Netil or 8-arm Netil were sacrificed on day 30. Serum and major organs were collected for biochemical analysis and hematoxylin and eosin (H&E) staining, respectively.

The systemic biosafety of 8-arm Netil was further assessed in healthy mice. Mice were randomly divided into control and 8-arm Netil groups. The 8-arm Netil group received intravenous injections of 8-arm Netil at 10 mg/kg at 1, 12, and 24 h, whereas control mice received an equal volume of saline through the same route. On day 30, serum and major organs were collected for H&E staining.

### Statistical analysis

Statistical analyses were performed using GraphPad Prism software (La Jolla, CA), and error bars indicate s.e.m. The differences between groups were assessed by either Student’s t-test (for simple two-sample comparison) or one-way analysis of variance (ANOVA) with Tukey’s post hoc test (for multiple comparisons). The Kaplan-Meier method was used to compare differences in survival rates. Pearson correlation analyses report the coefficient r, the number of data points (n), the two-sided P value, and the 95% confidence interval; given the limited number of conjugates available for several correlations, these analyses are described as exploratory. Exact P values are reported throughout rather than significance thresholds alone wherever feasible. P < 0.05 was considered statistically significant (95% confidence level).

## Acknowledgements

This work was supported by Columbia University (UR001585-01), National Natural Science Foundation of China (No. 32522051), National Key Research and Development Program of China (2024YFA1212000). The authors acknowledge the use of ChatGPT (OpenAI) for language editing and text polishing during manuscript preparation. All AI-assisted text was reviewed, edited, and approved by the authors, who take full responsibility for the content of the manuscript.

## Data Availability Statement

The data that support the findings of this study are available in the supplementary material of this article.

## ToC figure

In severe sepsis, bacterial infection promotes NET-associated chromatin accumulation, TLR4/TLR9-NF-κB activation, and cytokine amplification. A selective multivalent NET-associated chromatin neutralizer, 8-arm Netil, complexes extracellular DNA-histone structures while retaining antibacterial function, thereby reducing bacterial dissemination, neutralizing NET-driven inflammatory signaling, and promoting immune-homeostatic resolution with favorable systemic biosafety.

**ToC figure**

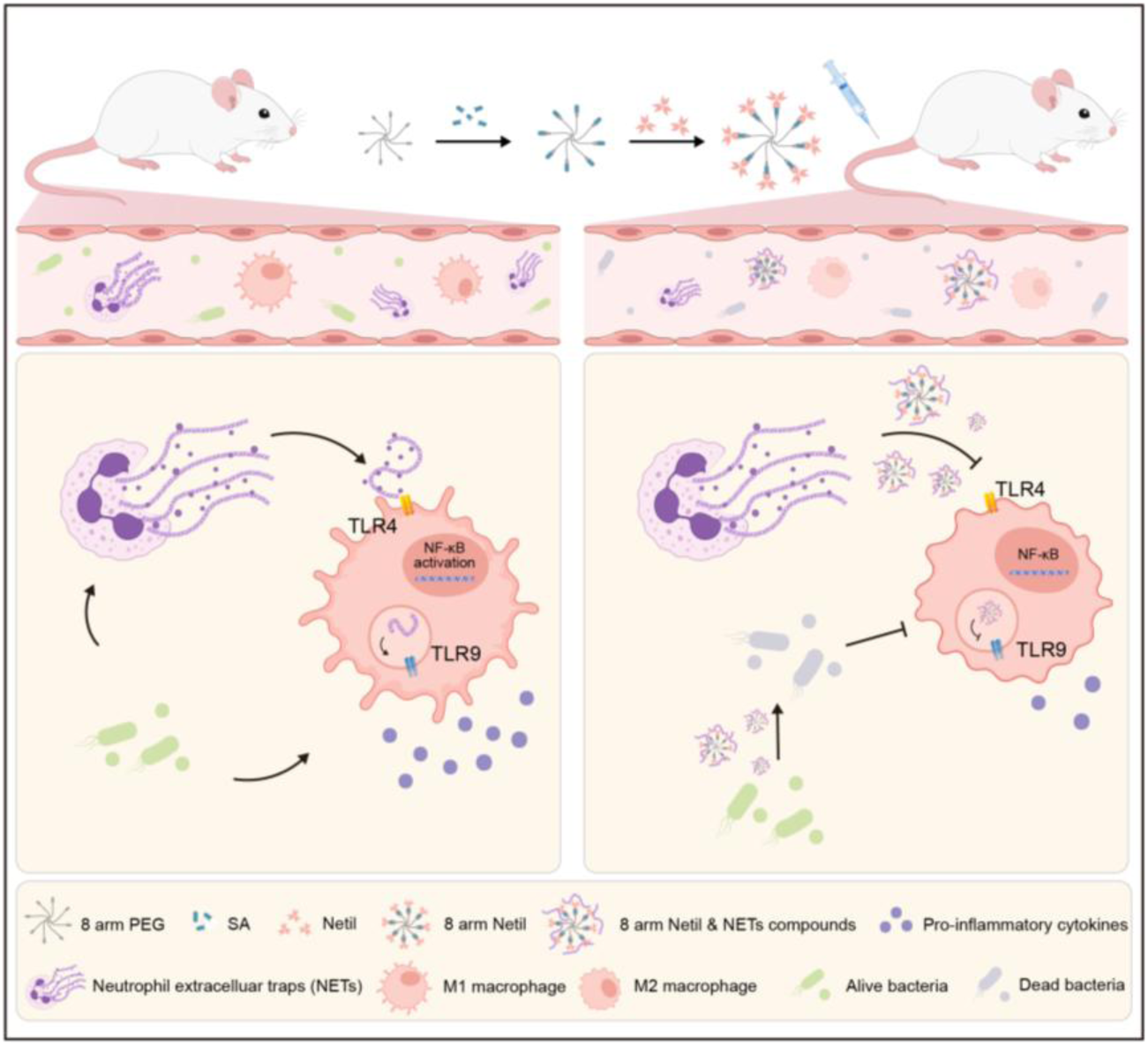

## References

[1] L. Evans, A. Rhodes, W. Alhazzani, et al., “Surviving Sepsis Campaign: International Guidelines for Management of Sepsis and Septic Shock 2021,” Crit. Care Med. 2021, 49, e1063–e1143.

[2] K. E. Rudd, S. C. Johnson, K. M. Agesa, et al., ”Global, regional, and national sepsis incidence and mortality, 1990-2017: analysis for the Global Burden of Disease Study,” Lancet 2020, 395, 200–211.

[3] M. Singer, C. S. Deutschman, C. W. Seymour, et al., “The Third International Consensus Definitions for Sepsis and Septic Shock (Sepsis-3),” JAMA 2016, 315, 801–810.

[4] J. E. Gotts, M. A. Matthay, ”Sepsis: pathophysiology and clinical management,” BMJ 2016, 353, i1585.

[5] J. M. Cavaillon, M. Singer, T. Skirecki, “Sepsis therapies: learning from 30 years of failure of translational research to propose new leads,” EMBO Mol. Med. 2020, 12, e10128.

[6] T. van der Poll, M. Shankar-Hari, W. J. Wiersinga, “The immunology of sepsis,” Immunity 2021, 54, 2450–2464.

[7] T. van der Poll, F. L. van de Veerdonk, B. P. Scicluna, M. G. Netea, “The immunopathology of sepsis and potential therapeutic targets,” Nat. Rev. Immunol. 2017, 17, 407–420.

[8] R. S. Hotchkiss, L. L. Moldawer, S. M. Opal, K. Reinhart, I. R. Turnbull, J. L. Vincent, ”Sepsis and septic shock,” Nat. Rev. Dis. Primers 2016, 2, 16045.

[9] J. L. Gross, R. Basu, C. J. Bradfield, et al., “Bactericidal antibiotic treatment induces damaging inflammation via TLR9 sensing of bacterial DNA,” Nat. Commun. 2024, 15, 10359.

[10] J. Dawulieti, M. Sun, Y. Zhao, et al., “Treatment of severe sepsis with nanoparticulate cell-free DNA scavengers,” Sci. Adv. 2020, 6, eaay7148.

[11] M. P. Fink, H. S. Warren, “Strategies to improve drug development for sepsis,” Nat. Rev. Drug Discov. 2014, 13, 741–758.

[12] V. Brinkmann, U. Reichard, C. Goosmann, et al., “Neutrophil extracellular traps kill bacteria,” Science 2004, 303, 1532–1535.

[13] V. Papayannopoulos, “Neutrophil extracellular traps in immunity and disease,” Nat. Rev. Immunol. 2018, 18, 134–147.

[14] S. R. Clark, A. C. Ma, S. A. Tavener, et al., “Platelet TLR4 activates neutrophil extracellular traps to ensnare bacteria in septic blood,” Nat. Med. 2007, 13, 463–469.

[15] A. Hakkim, B. G. Fürnrohr, K. Amann, et al., “Impairment of neutrophil extracellular trap degradation is associated with lupus nephritis,” Proc. Natl. Acad. Sci. U.S.A. 2010, 107, 9813–9818.

[16] A. Hidalgo, P. Libby, O. Soehnlein, I. V. Aramburu, V. Papayannopoulos, C. Silvestre-Roig, “Neutrophil extracellular traps: from physiology to pathology,” Cardiovasc. Res. 2022, 118, 2737–2753.

[17] G. Marsman, S. Zeerleder, B. M. Luken, “Extracellular histones, cell-free DNA, or nucleosomes: differences in immunostimulation,” Cell Death Dis. 2016, 7, e2518.

[18] J. Xu, X. Zhang, R. Pelayo, et al., “Extracellular histones are major mediators of death in sepsis,” Nat. Med. 2009, 15, 1318–1321.

[19] J. Xu, X. M. Zhang, M. Monestier, N. L. Esmon, C. T. Esmon, “Extracellular histones are mediators of death through TLR2 and TLR4 in mouse fatal liver injury,” J. Immunol. 2011, 187, 2626–2631.

[20] T. D. Tsourouktsoglou, A. Warnatsch, M. Ioannou, D. Hoving, Q. Wang, V. Papayannopoulos, “Histones, DNA, and citrullination promote neutrophil extracellular trap inflammation by regulating the localization and activation of TL R4,” Cell Rep. 2020, 31, 107602.

[21] J. Chen, T. Wang, X. O. Li, et al., “DNA of neutrophil extracellular traps promote NF-κB-dependent autoimmunity via cGAS/TLR9 in chronic obstructive pulmonary disease,” Signal Transduct. Target. Ther. 2024, 9, 162.

[22] C. H. O’Meara, L. A. Coupland, F. Kordbacheh, et al., ”Neutralizing the pathological effects of extracellular histones with small polyanions,” Nat. Commun. 2020, 11, 6408.

[23] H. Koide, A. Okishima, Y. Hoshino, et al., ”Synthetic hydrogel nanoparticles for sepsis therapy,” Nat. Commun. 2021, 12, 5552.

[24] P. X. Li, M. Li, M. R. Lindberg, M. J. Kennett, N. Xiong, Y. M. Wang, “PAD4 is essential for antibacterial innate immunity mediated by neutrophil extracellular traps,” J. Exp. Med. 2010, 207, 1853–1862.

[25] M. Mammen, S. K. Choi, G. M. Whitesides, ”Polyvalent interactions in biological systems: Implications for design and use of multivalent ligands and inhibitors,” Angew. Chem. Int. Ed. 1998, 37, 2754–2794.

[26] F. J. Martinez-Veracoechea, D. Frenkel, “Designing super selectivity in multivalent nano-particle binding,” Proc. Natl. Acad. Sci. U.S.A. 2011, 108, 10963–10968.

[27] B. Bruncsics, W. J. Errington, C. A. Sarkar, ” MVsim is a toolset for quantifying and designing multivalent interactions,” Nat. Commun. 2022, 13, 5029.

[28] S. Erlendsson, K. Teilum, “Binding Revisited-Avidity in Cellular Function and Signaling,” Front. Mol. Biosci. 2021, 7, 615565.

[29] A. J. Ruthenburg, H. Li, D. J. Patel, C. D. Allis, “Multivalent engagement of chromatin modifications by linked binding modules,” Nat. Rev. Mol. Cell Biol. 2007, 8, 983–994.

[30] C. A. Musselman, M. E. Lalonde, J. Côté, T. G. Kutateladze, “Perceiving the epigenetic landscape through histone readers,” Nat. Struct. Mol. Biol. 2012, 19, 1218–1227.

[31] J. Lee, J. W. Sohn, Y. Zhang, K. W. Leong, D. Pisetsky, B. A. Sullenger, “Nucleic acid-binding polymers as anti-inflammatory agents,” Proc. Natl. Acad. Sci. U.S.A. 2011, 108, 14055–14060.

[32] D. E. Brodersen, W. M. Clemons Jr., A. P. Carter, R. J. Morgan-Warren, B. T. Wimberly, V. Ramakrishnan, “The structural basis for the action of the antibiotics tetracycline, pactamycin, and hygromycin B on the 30S ribosomal subunit,” Cell 2000, 103, 1143–1154.

[33] Q. Vicens, E. Westhof, “Crystal structure of paromomycin docked into the eubacterial ribosomal decoding A site,” Structure 2001, 9, 647–658.

[34] B. François, R. J. M. Russell, J. B. Murray, et al., “Crystal structures of complexes between aminoglycosides and decoding A site oligonucleotides: role of the number of rings and positive charges in the specific binding leading to miscoding,” Nucleic Acids Res. 2005, 33, 5677–5690.

[35] M. Chittapragada, S. Roberts, Y. W. Ham, “Aminoglycosides: molecular insights on the recognition of RNA and aminoglycoside mimics,” *Perspect*. Med. Chem. 2009, 3, 21–37.

[36] D. Rittirsch, M. S. Huber-Lang, M. A. Flierl, P. A. Ward, ”Immunodesign of experimental sepsis by cecal ligation and puncture,” Nat. Protoc. 2009, 4, 31-36.

[37] C. Dufès, I. F. Uchegbu, A. G. Schätzlein, “Dendrimers in gene delivery,” Adv. Drug Deliv. Rev. 2005, 57, 2177–2202.

[38] G. L. Burn, T. Raisch, S. Tacke, et al., “Myeloperoxidase transforms chromatin into neutrophil extracellular traps,” Nature 2025, 647, 747–756.

[39] J. A. Buras, B. Holzmann, M. Sitkovsky, “Animal models of sepsis: Setting the stage,” Nat. Rev. Drug Discov. 2005, 4, 854–865.

[40] K. Buscher, H. Y. Wang, X. L. Zhang, et al., ”Protection from septic peritonitis by rapid neutrophil recruitment through omental high endothelial venules,” Nat. Commun. 2016, 7, 10828.

[41] A. Mai, M. Esposito, G. Sbardella, S. Massa, “A new facile and expeditious synthesis of N-hydroxy-N′-phenyloctanediamide, a potent inducer of terminal cytodifferentiation,” Org. Prep. Proced. Int. 2001, 33, 391–394.

[42] Y. L. Chen, F. M. Chen, X. H. He, et al., “Myeloperoxidase-mimetic nanozyme generates hypochlorous acid for phagosomal bacteria elimination,” Nano Today 2024, 54, 102137.

[43] A. L. Wishart, M. Swamydas, M. S. Lionakis, ”Isolation of Mouse Neutrophils,” Curr. Protoc. 2023, 3, e879.

[44] K. B. Zhang, C. Yang, C. X. Cheng, et al., ”Bioactive Injectable Hydrogel Dressings for Bacteria-Infected Diabetic Wound Healing: A “Pull-Push” Approach,” ACS Appl. Mater. Interfaces 2022, 14, 26404–26417.

[45] C. Yang, J. Dawulieti, K. B. Zhang, et al., “An Injectable Antibiotic Hydrogel that Scavenges Proinflammatory Factors for the Treatment of Severe Abdominal Trauma,” Adv. Funct. Mater. 2022, 32, 2111698.

[46] Y. He, C. X. Cheng, Y. H. Liu, et al., “Intravenous Senescent Erythrocyte Vaccination Modulates Adaptive Immunity and Splenic Complement Production,” ACS Nano 2024, 18, 470–482.

